# Translation Inhibitory Elements from Hox a3 and a11 mRNAs use uORFs for translation inhibition

**DOI:** 10.1101/2021.01.19.427285

**Authors:** Fatima Alghoul, Laure Schaeffer, Gilbert Eriani, Franck Martin

## Abstract

During embryogenesis, Hox mRNA translation is tightly regulated by a sophisticated molecular mechanism that combines two RNA regulons located in their 5’UTR. First, an Internal Ribosome Entry Site (IRES) enables cap-independent translation. The second regulon is a Translation Inhibitory Element or TIE, which ensures concomitant cap-dependent translation inhibition. In this study, we deciphered the molecular mechanisms of Hox a3 and a11 TIE elements. Both TIEs possess an upstream Open Reading Frame (uORF) that is critical to inhibit cap-dependent translation. However, the molecular mechanisms used are different. In TIE a3, we identify a uORF which inhibits cap-dependent translation and we show the requirement of the non-canonical initiation factor eIF2D for this process. The mode of action of TIE a11 is different, it also contains a uORF but it is a minimal uORF formed by an uAUG followed immediately by a stop codon, namely a ‘start-stop’. The a11 ‘start-stop’ sequence is located upstream of a highly stable stem loop structure which stalls the 80S ribosome and thereby inhibits cap-dependent translation of Hox a11 main ORF.

## Introduction

Gene expression constitutes an indispensable cellular process for which the genetic information encodes a functional product, mainly proteins. This process named translation initiates by a cap-dependent mechanism for most cellular mRNAs. It involves a large number of auxiliary proteins termed eukaryotic Initiation Factors (eIFs) which are required for the recruitment of the ribosomes on the mRNA (Hinnebusch, 2014; Merrick and Pavitt, 2018; Pelletier and Sonenberg, 2019; Shirokikh and Preiss, 2018). To ensure fine-tuning of translation, this step is highly regulated. However, several mRNA subclasses are translated by non-canonical mechanisms. For instance, this is the case for homeobox (Hox) mRNAs. Hox genes encode a family of proteins that constitutes transcription factors. Their main function is to orchestrate specific sequential transcription processes during embryonic development. A wealth of experimental data over the last three decades has led to the identification of many *cis*-regulatory elements that control Hox gene transcriptional patterns, thus giving deeper insights into the expression of Hox mRNAs (Alexander et al., 2009). In fact, the expression of the Drosophila genes Antp and Ubx have been suggested to be under translational control during embryonic development (Oh et al., 1992). More precisely, a subgroup of mRNAs produced from the Antp and Ubx loci contain functional Internal Ribosome Entry Sites (IRES) that allow their translation using a cap-independent mechanism. The IRES activity is modulated during development (Ye et al., 1997). More recently, the presence of other IRES elements in the 5’UTR of subsets of mice HoxA mRNAs (*Hoxa3, Hoxa4, Hoxa5, Hoxa9 and Hoxa11*) have been demonstrated (Leppek et al., 2020; Xue et al., 2015). Some of these IRESes require the presence of the ribosomal protein RpL38 in the ribosome to efficiently initiate translation, thereby explaining the tissue patterning defective phenotype observed with RpL38 knockout mouse (Kondrashov et al., 2011). The IRES activity in these Hox mRNAs was found to be critical for their appropriate expression. Upon the discovery of the IRES elements in Hox mRNAs, other RNA regulons termed Translational Inhibitory Elements (TIE) were also found in the 5’UTR of Hox mRNAs (Xue et al., 2015). TIEs are located upstream of the previously described IRES. According to the study by *Xue et al*., these elements efficiently inhibit canonical cap-dependent translation in subsets of HoxA mRNAs (*Hoxa3, Hoxa4, Hoxa9 and Hoxa11*) by an unknown mechanism (Xue et al., 2015). The action of TIE that ensures efficient blockage of cap-dependent translation, promotes IRES-mediated cap-independent translation. TIE and IRES act in synergy to ensure tightly regulated translation during organismal development. Indeed, it has been shown that Hox TIE elements ensure that Hox mRNAs are translated solely by their IRES element. Thereby, TIE elements represent the first example of specific RNA elements dedicated to inhibiting specifically cap-dependent translation in Hox mRNAs. However, the mechanism of action of these elements as well as their structural characterization are still unknown. In this study, we investigate the functional mode of action of TIE elements from a3 and a11 Hox mRNAs that were previously identified and characterized by *Xue et al*. (Xue et al., 2015). First, we determined their secondary structure by chemical probing assays and then, using cell-free translation extracts and *in vivo* assays, we deciphered their mode of action. Interestingly, the translation inhibitory mechanism that is mediated by a3 TIE element is radically distinct from the one used by a11. TIE a3 contains a uORF that is translated into a 9 KDa protein through the Hox a3 IRES and that requires the presence of the non-canonical translation initiation factor eIF2D. Indeed, eIF2D has been shown to be involved in diverse functions from translation initiation on specific mRNAs to reinitiation and recycling (Dmitriev et al., 2010; Skabkin et al., 2010; Weisser et al., 2017). On the contrary, TIE a11 contains an uAUG followed by a stop codon and a long stable hairpin. These three elements enable a highly efficient inhibition of cap-dependent translation by TIE a11 that is achieved through a start-stop stalling mechanism of an 80S ribosome.

## Materials and Methods

### Plasmids

For *in vitro* studies, murine TIE a3 (170 nts) and TIE a11 (216 nts) (sequences were kindly provided by Dr. Maria Barna, they were amplified from mouse E10.5–12.5 cDNA (Xue et al., 2015)) were placed upstream of 5’UTR of human β-globin (Accession number: KU350152) (50 nts) and *Renilla reniformis* Luciferase coding sequence (Accession number: M63501) (936 nts). These constructs were cloned in pUC19 vector in the *HindIII* site then used as a template for further PCR amplifications and site-directed mutagenesis.

For *in vivo* studies, we introduced an *Eco*RI restriction site upstream of hRLuc-neo fusion sequence in pmirGLO vector (Promega^®^) using Quick Change site-directed mutagenesis kit II XL (Thermo Fischer Scientific^®^). Subsequent cloning experiments of TIE a3, TIE a11 and their mutants were performed in pmirGLO vector at EcoRI site using NEBuilder^®^ *HiFi* DNA Assembly kit. All clones were checked by sequencing.

### Cell lines

Two cell lines were used for our *in vivo* assays: Human embryonic kidney cell line HEK293FT (ATCC^®^) and murine mesenchymal stem cell line C3H10T1/2 (clone 8, ATCC® CCL26). HEK293FT cells were cultured in Dulbecco’s modified Eagle medium (DMEM) with 2 mM of L-Glutamine and 10% Fetal Bovine Serum (FBS) supplemented with 100 units/ml of Penicillin/Streptomycin. Subcultures were performed after Trypsin-EDTA treatment for dissociation at sub-confluent conditions (70%-80%) 1:4 to 1:10 seeding at 2-4.10^4^ cells/cm^2^ according to manufacturer’s instructions. C3H10T1/2 cells were cultured in basal DMEM medium supplemented with 2 mM Glutamine, 1.5 g/L sodium bicarbonate and 10% FBS supplemented with 40 μg/ml Gentamicine. Subcultures were performed after Trypsin-EDTA treatment for dissociation at sub-confluent conditions (60%-70%). Seeding dilutions were performed at 2000 cells/cm^2^ one time per week.

### RNA transcription

Transcription templates were generated by PCR amplification from the plasmids pUC19-TIE. The amplified templates were used for *in vitro* transcription with recombinant T7 RNA polymerase in the presence of either m^7^G_ppp_G cap analog or non-functional cap analog A_ppp_G (New England Biolabs®). To check RNA integrity, an aliquot was mixed with Formamide Dye and loaded on 4% denaturing polyacrylamide gel. The RNA is visualized under UV light after ethidium bromide staining. To eliminate unincorporated nucleotides, the RNA sample was loaded on a gel filtration Sephadex G25-column (Pharmacia Fine Chemicals), proteins are then eliminated by phenol extraction and the RNA transcripts are precipitated with 0.25 M NaCl in ethanol. After centrifugation, RNA pellets were dried and resuspended in autoclaved milli-Q water. The concentration of purified RNA samples was determined by absorbance measurement at 260 nm.

### *In vitro* translation assays in cell-free translation extracts

*In vitro* translation was carried out using increasing concentrations of mRNA transcripts with self-made untreated Rabbit Reticulocyte Lysate (RRL), amino acid mixture containing all the amino-acids except methionine (1 mM of each), RNasin (Promega ®), 75mM KCl, 0.5 mM MgCl_2_, 3.8 mCi [^35^S] methionine and autoclaved milli-Q water. Reaction mixture was incubated at 30°C for 1 hr. Aliquots of translation mixture were analysed by SDS PAGE (10%) (Laemmli, 1970) and translation products were visualized by phosphor-imaging. *In vitro* translation assays with wheat germ extract (Promega®) were performed according to manufacturer’s instructions. *In vitro* translation assays with HeLa cell extract and drosophila S2 cell extract were performed as previously described (Thoma et al., 2004; Wakiyama et al., 2006).

### Chemical probing

#### Probing with DMS

Modification by Dimethyl Sulfate (DMS) was performed on 2 pmoles of each RNA (TIE a3 and TIE a11). The RNA is first incubated for 15 min in dimethylsufate (DMS) buffer (50 mM Na Cacodylate (pH 7.5), 5 mM MgCl_2_ and 100 mM KCl) and 1 μg of yeast total tRNA (Sigma-Aldrich®) and then modified with 1.25% DMS reagent (diluted with ethanol 100%) with 10 min incubation at 20°C and stopped on ice. Modified transcripts are precipitated with 0.25 M NaCl, 0.1 mg/ml glycogen in ethanol. RNA pellets were dried and resuspended in autoclaved milli-Q water. Modified nucleotides were detected by primer extension arrests that were quantified. The intensity of the RT stops is proportional to the reactivity for each nucleotide.

#### Probing with CMCT

Similarly, modification by 1-cyclohexyl-3-(2-morpholinoethyl) carbodiimide metho-p-toluene sulfonate (CMCT) was performed on 2 pmoles of each RNA (TIE a3 and TIE a11). Each RNA is incubated for 20 min in CMCT buffer (50 mM Na borate (pH 8.5); 5 mM MgCl_2_; 100 mM KCl) and 1 μg of yeast total tRNA. Then modifications were performed with 10.5 g/L CMCT reagent with 20 min incubation at 20°C and stopped on ice. Modified transcripts are precipitated with 0.25 M NaCl, 0.1 mg/ml glycogen in ethanol. RNA Pellets were dried and resuspended in autoclaved milli-Q water. Modified nucleotides were detected by primer extension arrests that were quantified. The intensity of the RT arrests is proportional to the reactivity for each nucleotide.

### Primer extension

Reverse transcription was carried out in a 20 μl-reaction with 2 pmoles of RNA and 0.9 pmoles of 5’ fluorescently labelled primers. We used fluorescent Vic and Ned primers (Thermo Fischer Scientific) of same sequence for all reverse transcription reactions which are complementary to the β-globin 5’UTR from nucleotides 6-37 : 5’-GGTTGCTAGTGAACACAGTTGTGTCAGAAGC-3’. First, the RNA was unfolded by a denaturation step at 95°C for 2 min. Then, fluorescent primers are annealed for 2 min at 65°C followed by incubation on ice for 2 min. Samples are incubated in a buffer containing 83 mM KCl, 56 mM Tris-HCl (pH 8.3), 0.56 mM each of the four deoxynucleotides (dNTP), 5.6 mM DTT and 3 mM MgCl_2_. Reverse transcription was performed with 1 unit of Avian Myoblastosis Virus (AMV) reverse transcriptase (Promega®) at 42°C for 2 min, 50°C for 30 min and finally 65°C for 5 min. In parallel, sequencing reactions were performed in similar conditions, but supplemented with 0.5 mM dideoxythymidine or or dideoxycitidine triphosphate (ddTTP or ddCTP) (protocol adapted from (Gross et al., 2017)). The synthesized cDNA were phenol–chloroform extracted and precipitated. After centrifugation, the cDNA pellets were washed, dried and resuspended in 10 μl deionized Hi-Di formamide (freshly prepared highly deionized formamide). Samples were loaded on a 96-well plate for sequencing on an Applied Biosystems 3130xl genetic analyzer. The resulting electropherograms were analyzed using QuSHAPE software (Karabiber et al., 2013), which aligns signal within and across capillaries, as well as to the dideoxy references of nucleotide at specific position and corrects for signal decay. Normalized reactivities range from 0 to 2, with 1.0–2.0 being the range of highly reactive positions. A preliminary secondary structure model was first initiated by mfold (Mathews et al., 2016) and then edited according to reactivity values.

### Sucrose gradient analysis

To analyse the assembly of ribosomal preinitiation complexes on the RNA of interest, the complexes were loaded on 7-47% sucrose gradients containing 5 mM MgCl_2_, 25 mM Tris-HCl (pH 7.5), 1 mM DTT and 50 mM KCl. We used the specific translation inhibitors GMP-PNP (4 mM), cycloheximide (1 mg/mL), geneticin (0.7 mM), hygromycin (0.5 mg/mL) and edeine (10 mM), they were added to the RRL with a mix containing the 20 amino acids at 1.5 mM each, RNasin (Promega®), 35 mM KCl and 0.24 mM or 2.4 mM MgCl_2_, prior to incubation with the 5’ capped radioactive mRNA of interest. The assembled pre-initiation complexes were formed by incubation in RRL at 30°C for 5 min. Then, 8 mM MgAc2 was added and one volume of 7% sucrose. Samples were then layered on the surface of 11-mL 7-47% sucrose gradient and centrifuged for 2h30 in an SW41 rotor at 37,000 rpm at 4°C. After centrifugation, the whole gradient is fractionated, and the mRNA is localized by measuring radioactivity in each collected fraction by Cerenkov counting in a scintillation counter.

### Mass spectrometry and data processing

Protein extracts were digested with sequencing-grade trypsin (Promega®) as previously described (Chicher et al., 2015; Prongidi-Fix et al., 2013). Peptide digests were analysed by nano LC-MS/MS and MS data were searched by the Mascot algorithm against the UniProtKB database from *Oryctolagus cuniculus* (rabbit). Identifications were validated with a protein False Discovery Rate (FDR) of less than 1% using a decoy database strategy. The total number of MS/MS fragmentation spectra was used to quantify each protein from three independent biological replicates. This spectral count was submitted to a negative-binomial test using an edge R GLM regression through the R-package. For each identified protein, an adjusted P-value corrected by Benjamini–Hochberg was calculated, as well as a protein foldchange (FC is the ratio of the average of spectral counts from a specific complex divided by the average of spectral counts from a reference protein complex). The results are presented in a volcano plot using protein log_2_ FC and their corresponding adjusted log_10_ P-values. The proteins that are up-regulated in each condition are shown in red (TIE a3 versus β-globin mRNA, TIE a11 versus β-globin mRNA and TIE a3 versus TIE a11).

### *In vivo* luciferase assay

For *in vivo* luciferase assay, HEK293T cells and C3H10T1/2 cells were transfected in 6-well plates with various constructs of pmirGlo vector (Promega®). Transfection was performed using Turbofect transfection reagent (Thermo Fischer scientific®) according to manufacturer’s instructions. Cells were collected 24 hrs post-transfection. Luciferase assay was performed using Dual-Glo luciferase kit (Promega®) according to manufacturer’s instructions. Firefly Luciferase activities were measured to monitor transfection efficiency in order to normalize Renilla luciferase activities for each construct.

### Co-transfection assay of siRNAs and reporter plasmid in HEK293T cells

HEK293FT cells were used to test the effect of inhibition of eIF2D knock-down, eIF4E knock down and a non-target siRNA pool as a negative control on Renilla luciferase expression (*hRluc-neo*) in pmirGLO vectors. We used ON-TARGETplus human siRNAs against eIF2D (catalog number L-003680-01-00), siRNA against eIF4E (catalog number J-003884-10-00) and non-target pool of siRNAs (catalog number D-001810-10-05) purchased from Horizon Discovery. HEK293T cells were cultured according to manufacturer’s instructions (ATCC®) for 24 hr and used for co-transfection by siRNAs and reporter plasmid. We used 2×10^5^ cells in 1 ml of culture medium without antibiotics. Upon reaching 70% confluence, cells were transfected by 5 pmoles of siRNAs in different wells with 500 ng of reporter plasmid. Transfections were performed with Lipofectamine 2000 (Invitrogen®) according to manufacturer’s instructions. After 48h, cells were washed twice by phosphate buffered saline (PBS1X) and incubated with Passive Lysis Buffer (PLB) 1X for 15 min. Luciferase assay was performed according to manufacturer’s instructions. Protein concentration was measured by Bradford’s assay. The impact of silencing on TIE-mediated translation inhibition was measured by luciferase assay according to manufacturer’s instructions as previously mentioned.

### Western Blot against eIF2D, eIF4E and GAPDH

The silencing efficiency was quantified by western blots using rabbit polyclonal anti-eIF2D antibody (12840-1-AP) from Proteintech, mouse monoclonal anti-eIF4E antibody (sc-9976) and mouse polyclonal anti-GAPDH antibody (sc-1377179) from Santa Cruz Biotechnology. For that, 20 μg of protein extracts were loaded on 10% polyacrylamide SDS PAGE. After migration, proteins were transferred to an Immobilin-P membrane (Millipore®) at 10 Volts for 1 h in a semi-dry apparatus (Trans-Blot® SD) on a PVDF membrane (PolyVinyliDene Fluoride) that had been previously activated with 100% methanol for few seconds and a transfer buffer pH 8 (25 mM Tris; 200 mM glycine; 20% ethanol). After transfer, the membrane was saturated for 2h by blocking buffer (5% milk, 0.05% Tween-20, PBS 1X). Primary antibodies were added at dilutions recommended by the manufacturers in blocking buffer, the membranes were incubated overnight at 4°C. Then, the membranes were washed three times by PBST (PBS 1X; 0.05% Tween-20) to remove the excess of primary antibodies. Then, membranes were incubated with secondary HRP-conjugated antibody for 1h at room temperature followed by three washing steps. The signal produced by reaction between HRP and ECL (Kit ECL Plus Western Blotting Detection System, GE Healthcare®) was detected by chemiluminescence using imaging Chemidoc (Biorad®).

### Ribosome toe printing assay

Toe printing assay was adapted from previously established protocols (Martin et al., 2011, 2016). Briefly, RRL were incubated for 5 min at 30°C then 10 min on ice with buffer containing 1U/μl of recombinant RNasin (Promega®), 75 mM KCl 0.5 mM MgCl_2_, and 1.3 mM of puromycin prior to initiation complex assembly. Then, the pre-initiation complexes are formed by incubation with 500 nM of the RNA of interest in the presence of specific inhibitors such as cycloheximide (1 mg/ml) or GMP-PNP (4 mM) for 5 min at 30°C and then 20 min on ice. Then, the pre-initiation complexes were complemented with one volume of ice-cold buffer A containing 20 mM Tris-HCl (pH 7.5), 100 mM KAc, 2.5 mM MgAc_2_, 2 mM DTT, 1 mM ATP and 0.25 mM spermidine and placed on ice. In order to separate ribosomal complexes from the non-ribosomal fraction, samples were centrifuged at 88,000 rpm in S100AT3 rotor (Sorvall-Hitachi) at 4°C for 1h. After centrifugation, the pellets containing the pre-initiation complexes were resuspended in 30 μl of ice-cold buffer A and incubated with 5’ radioactively labelled DNA oligonucleotide complementary to nts 22-51 of renilla luciferase coding sequence for 3 min at 30°C. Then, 1 μl of a 320 mM Mg(Ac)_2_, 4 μl of a dNTP mixture (containing 5 mM of dATP, dGTP, dTTP and dCTP), 10 units of recombinant RNAsin (Promega®) and 1 unit/μl Avian Myoblastosis virus reverse transcriptase (Promega®) were added and incubated for 1h at 30°C. The synthesized cDNAs were analysed on 8% PAGE next to sequencing ladders.

## Results

### TIE-mediated inhibition recapitulated in RRL

To dissect the mechanism of TIE-mediated inhibition, we first set-up an efficient cell-free assay. We used RRL and performed *in vitro* experiments using several reporter constructs. To analyse their translation efficiency, capped mRNAs were translated in RRL in the presence of [^35^S]-methionine to verify the size of the expected synthesized protein and luciferase activity was measured to determine their translation efficiency. In order to standardize the measured translation activities, we inserted the TIE elements identified and characterized by *Xue et al*. (Xue et al., 2015), in murine Hox a3 and a11 mRNAs upstream of the well-characterized human β-globin 5’UTR (**Fig. 1A**). The human β-globin allows translation strictly by a cap-dependent mechanism (Fletcher et al., 1990). Therefore, in order to evaluate the inhibition due to TIEs, we used a reporter mRNA containing the sole human β-globin 5’UTR as a reference. First, we performed translation experiments with increasing amounts of mRNA and determined that with concentrations higher than 100 nM, the translation yield reached a plateau (**Fig. S1A**). In order to avoid any titration effect due to excess of mRNA, we performed all translation assays at an mRNA concentration of 50 nM, which enables sub-saturating conditions. TIE a3 and a11 efficiently promote a translation inhibition by 79% and 91% respectively (**Fig. 1A**). Therefore, our cell-free translation assay using RRL efficiently recapitulates the previously described TIE-mediated translation inhibition (Xue et al., 2015). Likewise, TIE-mediated inhibition was efficiently recapitulated in other cell-free translation systems from other organisms like Wheat Germ Extracts, drosophila S2 cell extracts and human HeLa cell extracts (**Fig. S1B**). Moreover, the same constructs were introduced in a plasmid that allowed us to monitor translation inhibition *in vivo*. We tested two cell lines that have been shown to express Hox genes, namely human HEK293T and murine mesenchymal stem cell line C3H10T1/2 (Phinney et al., 2005). In these cells, translation inhibition by TIE a3 and a11 is 98% and 88% respectively (**Fig. S1C**). Next, to define the minimal active domains of a3 and a11, sequential 5’ deletions were performed. In each experiment, translation for each construct was compared to the control (w/o TIE) (**Fig. 1B**). First, we tested large deletions in order to roughly determine the functional domain (**Fig. S1D**), then more precise deletions to map exactly the 5’ end of functional domains for a3 and a11 (**Fig. 1B**). Deletion of 68 nts in TIE a3 does not affect inhibition (81%). A further deletion of 113 nts completely abolishes the translation inhibition of TIE a3 (10%). Therefore, the minimal a3 element encompasses nucleotides 68 to 170. Concerning TIE a11, the situation is not as clear.

**Figure 1:**
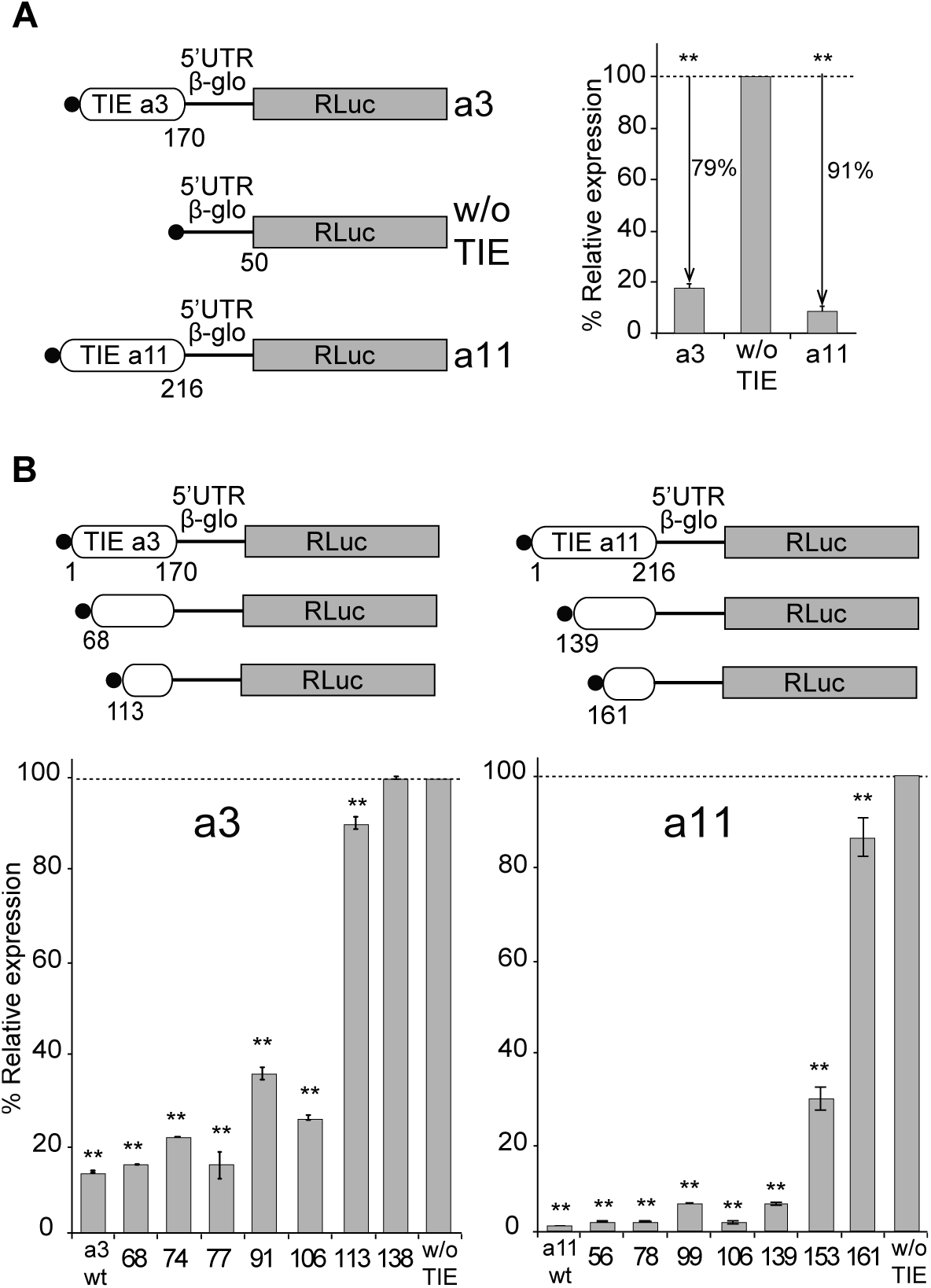
TIE-mediated inhibition is recapitulated in RRL and does not require full length TIEs. **A.** Three capped mRNAs were used to test TIE-mediated inhibition *in vitro*. TIE a3 and TIE a11 were placed upstream of the 5’UTR of β-globin and the Renilla luciferase coding sequence. Translation assays were performed *in vitro* using RRL at an mRNA concentration of 50 nM, which enables sub-saturating conditions. The relative expression of Luciferase protein reflects the efficiency of translation inhibition by TIE a3 and TIE a11. Values were normalized to that of the control (w/o TIE) which corresponds to normal expression without inhibition and was set to 100%. **p<0.01 (t-test as compared to w/o TIE) n=3. Experiments were performed in triplicates. **B.** Sequential deletions in the 5’ extremity of TIE a3 and TIE a11 constructs were performed to assay their effect on translation. Values of translation expression were normalized to that of the control (w/o TIE). Experiments were performed in triplicates. The percentages of inhibition for each TIE are indicated in the histogram. **p<0.01 (t-test as compared to w/o TIE) n=3. Experiments were performed in triplicates.

In fact, the inhibition is only partially reduced (70%) when 153 nts are deleted and completely lost when 161 nts are deleted. In this case, the deletions indicate that elements essential for translation inhibition are most likely located between residues 139 to 216. In conclusion, these experiments allowed the localisation of essential RNA elements required for a3 and a11 RNA regulons that are required to retain their full inhibitory function.

### TIE elements have distinct secondary structures

To gain further insights into the structural and functional properties of TIE elements, we performed chemical probing using DMS and CMCT reagents for both TIE a3 and TIE a11 (**Fig. 2**). Since modifications were performed in triplicates, the average of reactivity was calculated for each nucleotide at a specific position (**Fig. S2**). With this reactivity, we built a secondary structural model for both TIE a3 and a11. TIE a3 contains a 5’ proximal stem-loop and another bigger structure comprising two-way junctions (**Fig. 2A**). For TIE a11, the higher GC content (64%) than TIE a3 (45%) suggests more stable structure. Our probing experiments confirmed this statement. It comprises four stem-loops and a three-way junction structure (**Fig. 2B**). To further validate our models, the secondary structures of both TIEs were also probed in the frame of the full-length Hox 5’UTR which contains the IRES element downstream of the TIE. The reactivities for each nucleotide obtained with isolated TIEs and with TIEs embedded in full-length 5’UTR were highly similar suggesting that the TIEs do fold independently from the IRES (**Fig. S2**). Using these two models, we wanted to further characterize the structure-function relationships of both TIE elements.

**Figure 2:**
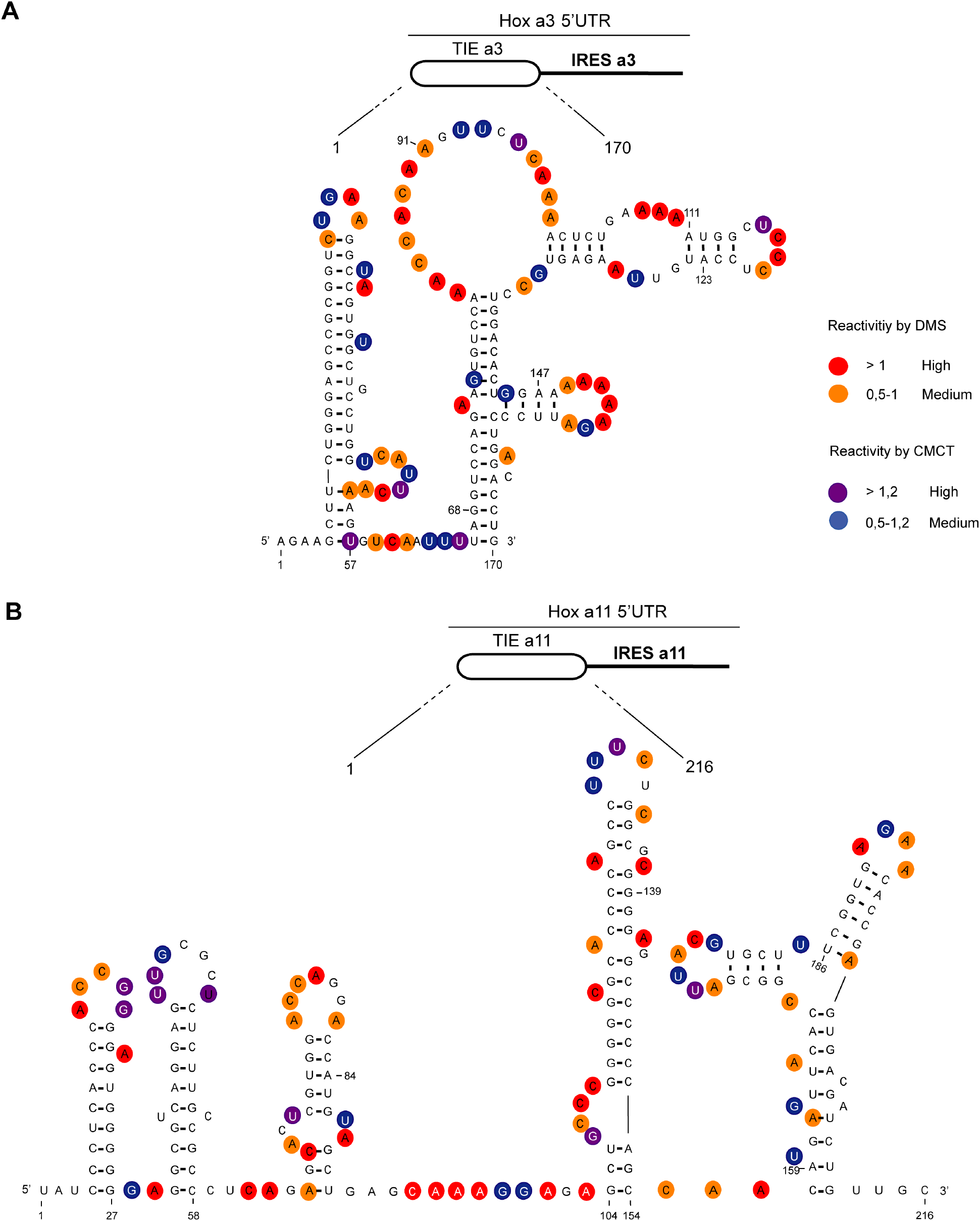
the secondary structural models of TIE a3 and TIE a11 reveal distinct structures. The structures of (A) TIE a3 (170 nucleotides) and (B) TIE a11 (216 nucleotides) were obtained by chemical probing using base-specific reagents, DMS and CMCT. After modifications, reverse transcription was performed using fluorescently labelled primers to determine the position of modified nucleotides. Experiments were performed in triplicates. Reactivities are shown as average reactivity from three independent experiments. A representation of reactivities is assigned as colour code depending on a range of values as shown in the figure legend on the right. Reactivity values for each nucleotide with corresponding standard deviations are shown in supplementary figures (S2A, S2B, S2C, and S2D).

### TIE a3 contains an upstream ORF that inhibits translation

The 5’ truncations experiments allowed us to pinpoint critical elements required for translation inhibition. Interestingly, the minimal a3 contains putative uORFs starting from two putative uAUGs at positions 111-113 and 123-125 respectively (**Fig. 3A**). In order to test their implication in the translation inhibition, both uAUGs were mutated independently into UAC thereby eliminating any possibility of AUG-like codon recognition. Interestingly, the mutation of AUG111 completely abolishes the inhibition by TIE a3 thereby confirming its implication. On the contrary, this is not the case for AUG123 (**Fig. 3A**). This is in good agreement with the fact that AUG111 has an optimal Kozak sequence (A at −3 and G at +4), thereby the ribosome initially recognizes it during scanning along the 5’UTR (Kozak, 1986). To further confirm the assembly of the ribosome on uAUG111, we mutated it into AUG-like codons. We tested previously described AUG-like codons such as AUU, CUG, GUG and ACG. None of the tested AUG-likes are used for translation inhibition indicating that TIE a3 requires a genuine AUG start codon (**Fig. S3A**). Since the AUG111 is involved in a stem, it is possible that the absence of inhibition with the AUG-likes is actually due to the disruption of the stem. To rule out this possibility, we mutated the AUG into CUG and GUG and inserted simultaneously the compensatory mutations that enable the formation of the stem with the mutated CUG and GUG codons. In both mutants, the inhibition was fully destroyed indicating that efficient inhibition requires a genuine AUG start codon independently of the secondary structure (**Fig. S3A**). Next, we confirmed the use of uAUG111 by toe printing assay. As expected, a canonical +16 reverse transcription arrest from uAUG111 was clearly detected with GMP-PNP and less efficiently with cycloheximide. Accordingly, when the uAUG is mutated to UAC, the +16 toe-print disappears (**Fig. 3B**). Altogether, this data confirm that a pre-initiation complex efficiently assembles on the uAUG111. We then wondered whether the ribosome assembled on this AUG codon proceeds to translation elongation resulting in a polypeptide synthesis encoded by the uORF. By performing sucrose gradient analysis, we detected polysome formation suggesting that efficient translation is occurring from uAUG111 (**Fig. S3B**). Accordingly, mutation of uAUG111 drastically reduces the amount of polysomes and the formation of 48S complexes in the presence of GMP-PNP thereby corroborating that the translation is starting on uAUG111 (**Fig. S3B**). In Hox a3 5’UTR, the uORF starting from uAUG111 is extending through the full IRES. To check further whether the uORF is indeed translated through the full length Hox a3 5’UTR, we first deleted a single nucleotide (G333) to change the frame and thereby producing a fusion protein formed by the peptide produced from the uORF and Renilla Luciferase. Indeed, with this single frameshifting point mutation, we could detect a longer protein demonstrating that the pre-initiation complex assembled on the uAUG111 indeed proceeds to translation elongation and is efficiently translating through the full length 5’UTR of Hox a3 mRNA (**Fig. 4A**). We also verified the translation from uAUG111 with our reporter constructs containing only the TIE a3 element. Likewise, the insertion of a single frameshifting nucleotide (A220) allows the detection of a longer fusion protein (**Fig. S4A**). Remarkably, with a double mutant that combines the mutation of the uAUG111 to ACG with the (A220) insertion, a mixture of two proteins is detected, the fusion protein and the more abundant Renilla Luciferase protein. Indeed, this confirms that the ribosome requires the uAUG111 but still able to recognize, although less efficiently, the AUG-like ACG as a start codon (**Fig. S4A**). Similarily, when we use the native full-length hox a3 mRNA, we detect the Hox a3 uORF protein of 9 KDa size (**Fig. S4B**). Altogether, our cell-free translation assays demonstrate that Hox TIE a3 translation is achieved by translation of a uORF that starts at AUG111 codon that extends through the whole Hox a3 5’UTR. To confirm these results *in vivo*, we generated reporter constructs in the plasmid pmirGlo that contains TIE elements upstream of Renilla luciferase (hRluc). Values were normalized to that of control enhanced Firefly luciferase (luc2) to calculate the luciferase activity for each report **(Fig. 4B)**. As expected, wild type TIE a3 blocks translation very efficiently in both HEK293T and C3H10T1/2 cells. When the uAUG111 is mutated to UAC, the inhibition is significantly affected in both HEK293T and C3H10T1/2 cells at respectively 57% and 45% compared to wild type TIE a3 (**Fig. 4B**). Although the inhibition is not fully abolished, these experiments confirmed that the codon AUG111 is critical for efficient translation inhibition in Hox a3 *in vivo*.

**Figure 3:**
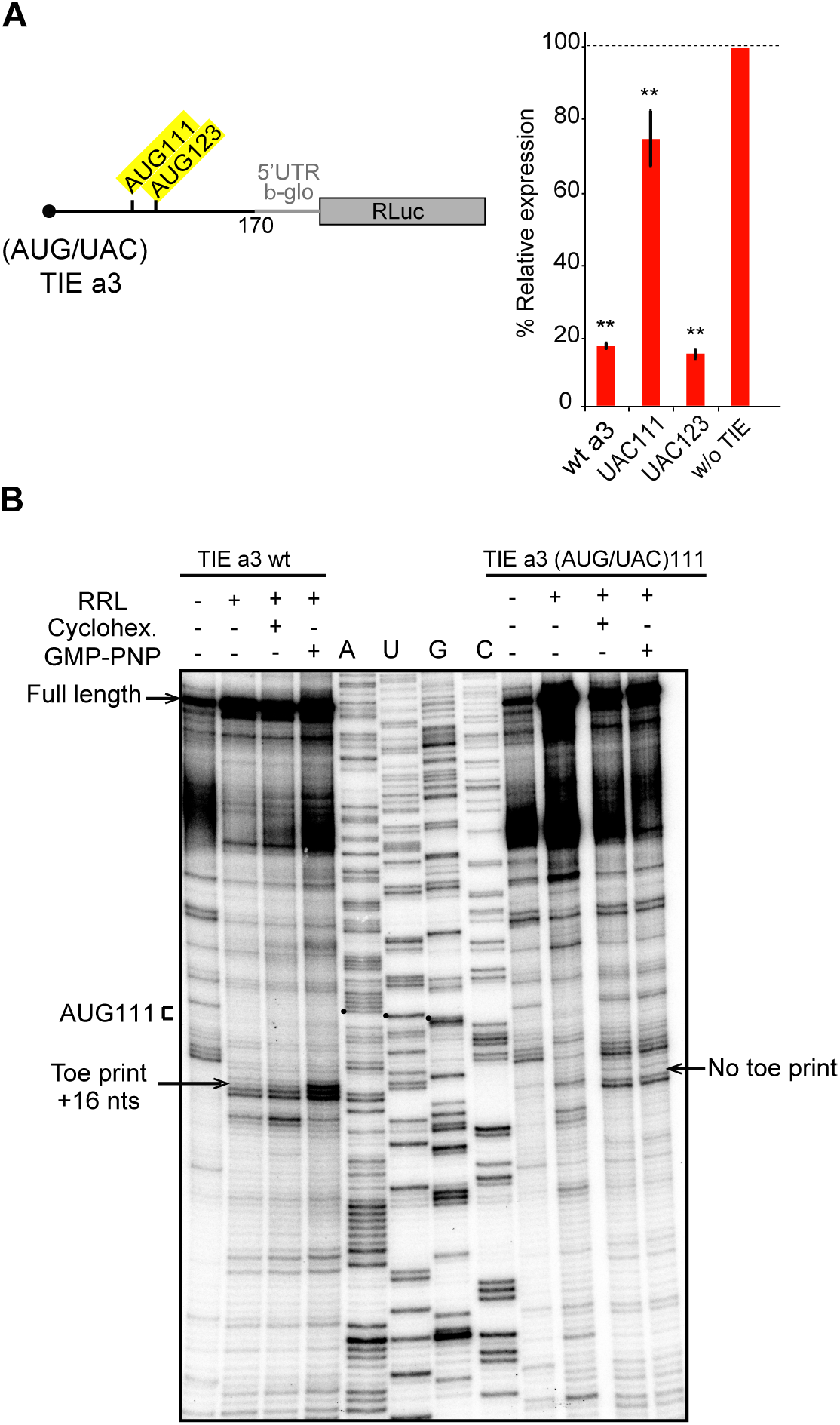
upstream AUG111 in TIE a3 is essential for inhibition. **(A)** Substitution mutations in uAUG111and uAUG123 to UAC in TIE a3 were performed. Constructs with the corresponding mutations were translated in RRL and luciferase assay was performed to evaluate the effect of mutation on translation efficiency as previously described. **p<0.01 (t-test as compared to construct w/o TIE). n=3. Experiments were performed in triplicates **(B)** Toe printing analysis of ribosomal assembly on two mRNAs, TIE a3 Wt and the mutant of upstream (AUG/UAC)111. Initiation complexes were assembled in RRL extracts in the absence or presence of translation inhibitors: cycloheximide and GMP-PNP. Reaction samples were separated on 8% denaturing PAGE together with the appropriate sequencing ladder. Toe-print positions were counted starting on the A+1 of the AUG codon at +16 position. Full-length cDNAs are indicated by an arrow at the top of the gel.

**Figure 4:**
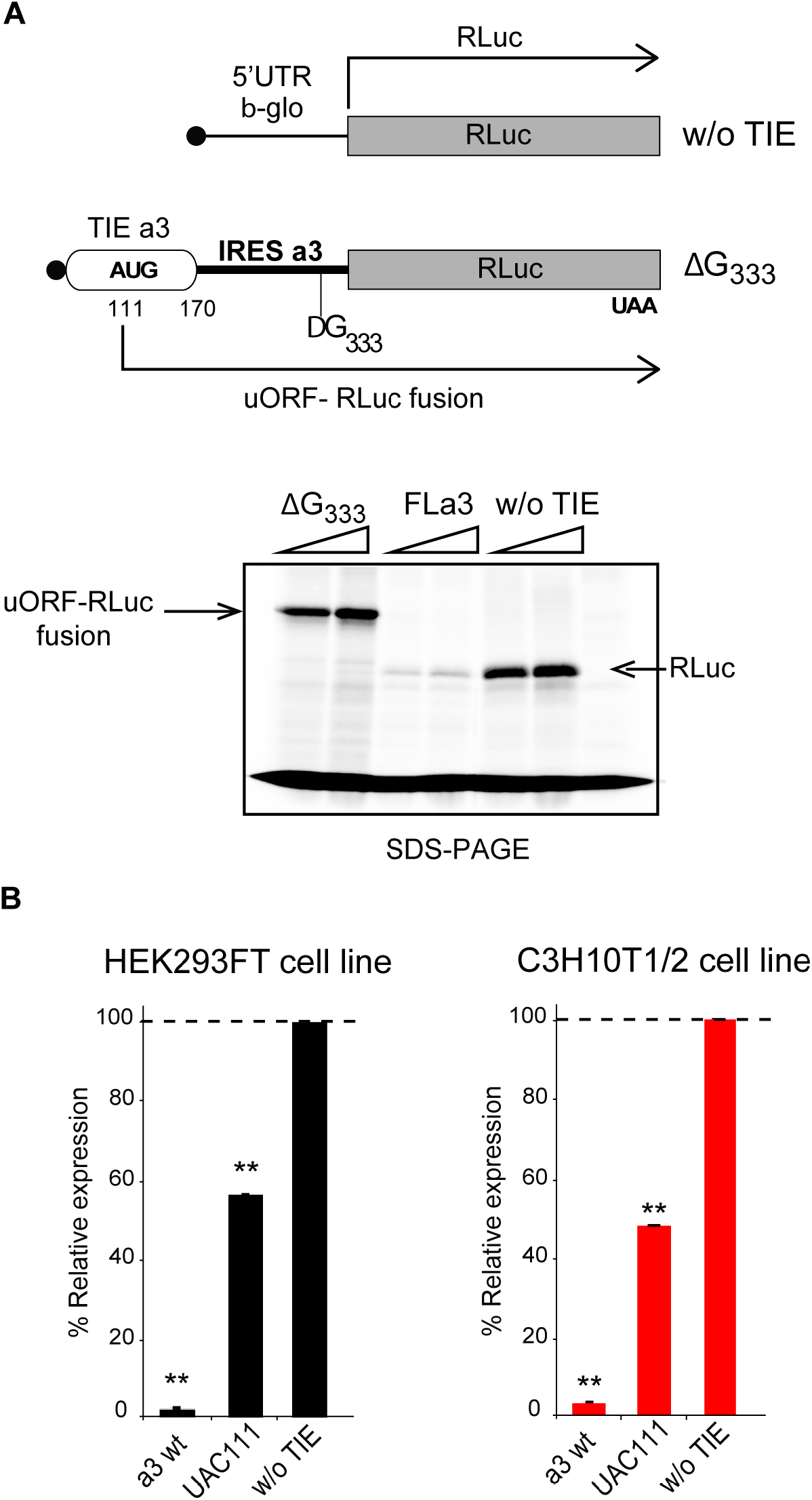
the uAUG111 in TIE a3 is translated through 5’UTR of Hox a3. **(A)** Three transcripts were used for this experiment: full length 5’UTR of Hox a3, a deletion mutant at nucleotide G333 in IRES a3 and a control transcript without TIE. To test the translation of uORF in TIE a3 starting form uAUG111, a deletion of G in IRES a3 at position 333 was performed to create a longer uORF that is in the same frame as the ORF of Renilla luciferase to create an N-terminally extended luciferase. Transcripts were translated *in vitro* in RRL and products were loaded on 10% SDS-PAGE in the presence of ^35^S-Methionine **(B)** *In vivo* luciferase assays in two embryonic cells lines; HEK293FT (left) and C3H10T1/2 (right). Reporter constructs in pmirGlo containing TIE a3, u(AUG/UAC)111, and without TIE, were transfected in the two indicated cell lines. Renilla luciferase expression was normalized to the control (w/o TIE), which was set to 100%. **p<0.01 (t-test as compared to empty plasmid (w/o TIE). Experiments were performed in triplicates n=3

### TIE a11-mediated inhibition is mediated by a stalled 80S ribosome

We next asked whether TIE a11 has a similar inhibitory mechanism. Deletion experiments suggested that critical elements were located in the region 139 to 216. We also found that TIE a11 contains two putative upstream AUGs at positions 84-86 and 159-161. Mutations of both AUG84 and AUG159 had no impact on translation inhibition (**Fig. 5A**). According to our 2D model, a long GC-rich stable stem loop (SL) structure (ΔG = −25.00 kcal/mol) spans nucleotides 104-154 (**Fig. 2B**). This long hairpin comprises sixteen G-C base pairs that can putatively interfere with the progression of a pre-initiation scanning complex. To test the inhibitory efficiency of this stem-loop, we transplanted it in a strictly cap-dependent reporter mRNA containing the 5’UTR of β-globin upstream of the luciferase coding sequence. The a11 SL was inserted in the middle of the 5’UTR, in this construct the first 25 nucleotides from the 5’ proximity are unfolded thereby ensuring proper access to the 5’ cap. Interestingly, the translation of this mRNA was significantly abolished showing that this stem loop on its own is sufficient to inhibit cap-dependent translation when transplanted in another mRNA (**Fig. S5**). Strikingly, in TIE a11, uAUG84 codon is located 19 nts upstream of the inhibiting SL. This distance is compatible with the assembly of a pre-initiation complex on uAUG84 without clashing with the SL. Moreover, the distance between the uAUG and the SL is optimal for favouring AUG recognition by scanning arrest of the pre-initiation complex forced by the SL. Importantly, the uAUG84 is immediately followed by a stop codon UAG87. The sequence context of the uAUG84 (A at −3 and U at +4) is suboptimal compared to the consensus Kozak sequence. This unique combination of start-stop codon upstream of stem loop structure raised the question of whether the ribosome is forced to recognize this uAUG despite a suboptimal Kozak context. To address this hypothesis, we mutated the stop codon UAG87 to UGG thereby creating a uORF. When the stop is mutated, a small peptide is produced from the translation of uAUG84 through 5’UTR, which we called a11 uORF (**Fig. 5B**). This experiment demonstrates that uAUG84 is indeed efficiently used as a start codon despite its sub-optimal sequence context. Unfortunately, the presence of the highly stable SL is not compatible with the toe-printing assay. Indeed, premature RT arrests occur due to the presence of the highly stable SL, rendering the toe-printing assay on TIE a11 impossible. Therefore, to further confirm that the ribosome is efficiently assembled on uAUG84, we performed sucrose gradient analysis with radiolabelled mRNAs. With wild type TIE a11, an 80S complex efficiently accumulates in the presence of cycloheximide as expected. However, an 80S complex is also detected without inhibitor indicating that the 80S complex is in fact a stalled ribosome (**Fig. 5C**). The mutation of uAUG84 to UAC drastically reduces the amount of 80S complex; in contrast, mutation of uAUG159 to UAC does not affect 80S accumulation. This further confirms that the stalled 80S complex is indeed assembled on the uAUG84. Altogether, this data show that a stalled ribosome is indeed assembled on uAUG84 and the stalling is caused by the synergistic effect of a stop codon next to the AUG and a stable SL downstream of the start-stop of a11.

**Figure 5:**
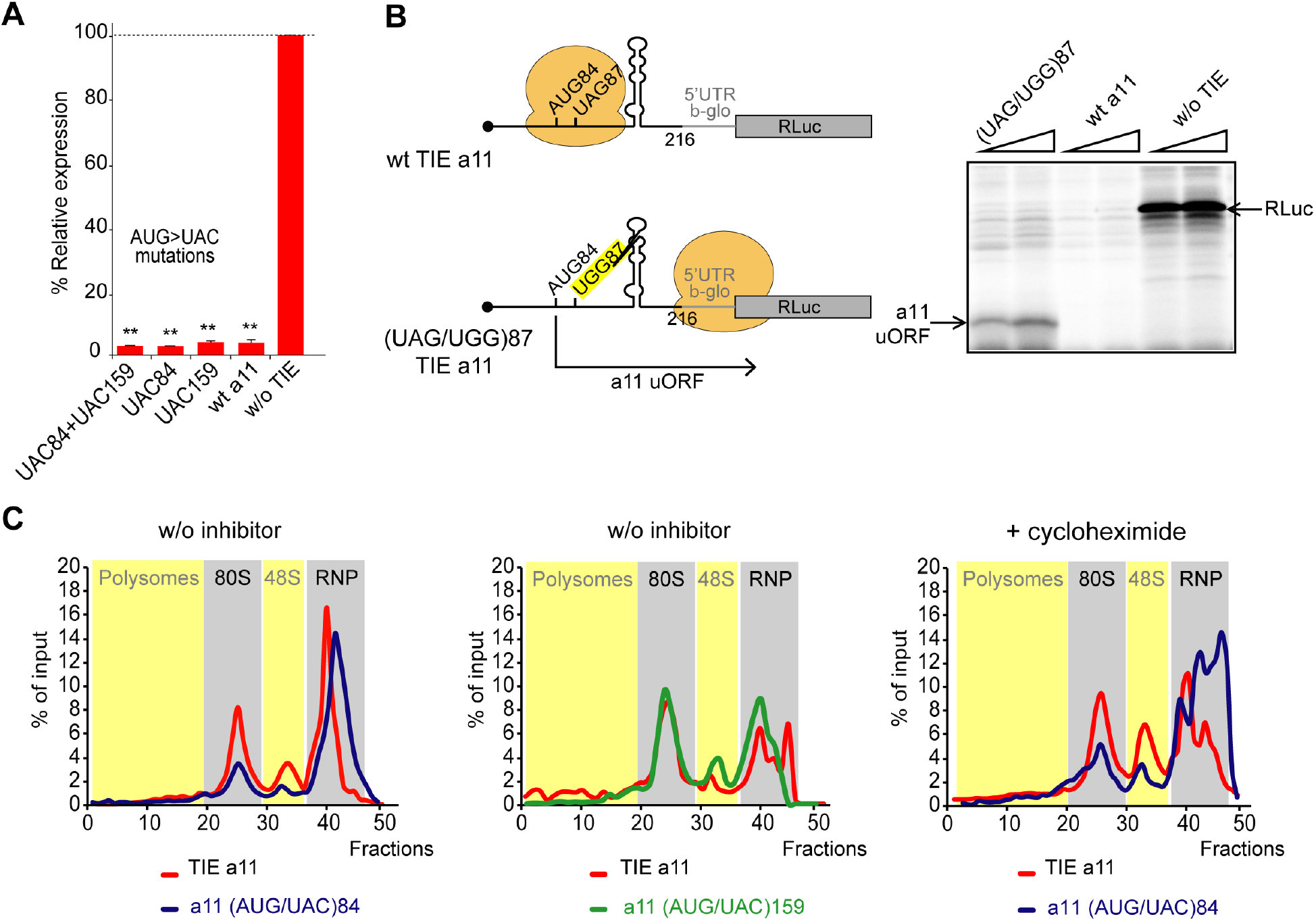
a start-stop uORF in TIE a11 stalls an 80S upstream of a highly stable structure. **(A)** Mutational analysis of uAUGs in TIE a11. Three transcripts with AUG/UAC mutations were used: M1: (AUG/UAC)84 + (AUG/UAC)159, M2: (AUG/UAC)84 and M3: AUG/UAC159. Transcripts were translated in RRL at 50 nM concentrations and the luciferase expression was normalized to the control (w/o TIE) as previously described. **p<0.01 (t-test as compared to construct w/o TIE). n=3. Experiments were performed in triplicates**. (B)** A single substitution (A/G)88 mutation destroys the stop codon UAG87 to UGG in TIE a11, TIE a11 wt and control (w/o TIE) were also translated as references in RRL. Translation products were loaded on 15% SDS-PAGE. **(C)** Ribosomal pre-initiation complexes were assembled and analysed on 7-47% sucrose gradient with [alpha-^32^P]GTP-radiolabeled TIE a11 as well as the two mutants of uAUG/UAC at the previously indicated positions in the absence or presence of cycloheximide. Heavy fractions correspond to polysomes and lighter fractions correspond to free RNPs.

### Mass spectrometry analysis pre-initiation complexes programmed by TIE a3 and a11

To further characterize the two different modes of action employed by a3 and a11, we identified the factors specifically acting in such mechanisms. For that, we purified preinitiation complexes programmed by TIE a3 and a11 suitable for Mass Spectrometry analysis. Briefly, ribosomes were assembled on chimeric biotinylated mRNA-DNA molecules and immobilized on streptavidin-coated beads after incubation with RRL in the presence of cycloheximide or GMP-PNP. Complexes were then eluted by DNase treatment as previously described (Chicher et al., 2015; Prongidi-Fix et al., 2013) and analysed by Mass Spectrometry. Three different mRNA constructs were used, TIE a3 upstream of 5’UTR β-globin, TIE a11 upstream of 5’UTR β-globin, and 5’UTR β-globin as a negative control. Comparison between the three mRNAs blocked with cycloheximide enabled the identification of specific factors interacting with each RNA (**Fig. 6**). Interestingly, for TIE a3, we identified a set of translation-related proteins. Among the strongest hits, we found eIF2D, a non-canonical GTP-independent initiation factor which has been shown to be involved in the initiation step on specific mRNAs (Dmitriev et al., 2010; Vaidya et al., 2017; Vasudevan et al., 2020), reinitiation on main ORF (Ahmed et al., 2018; Weisser et al., 2017) and recycling after translation termination (Skabkin et al., 2010, 2013; Young et al., 2018). Another interesting hit is methionine aminopeptidase MetAP1 which removes N-terminal methionine from nascent proteins in a co-translational manner (Varland et al., 2015). Other factors that are linked to translation have been selected such as the scanning factor eIF1A and its isoform eIF1A-X, eIF3a, Arginyl- and Leucyl-tRNA synthetases QARS and LARS, DEAD-box helicases DHX36 and DDX39B, elongation factor HBS1L and RpL38 ribosomal protein. For TIE a11, we identified different translation-related proteins among which are ASAP1, a GTPase activator protein, MetAP1, RpL38, Valyl-tRNA synthetase and eIF3j, another subunit of initiation factor eIF3 usually dissociating at early stages of initiation to allow mRNA entry (Aylett et al., 2015; Young and Guydosh, 2019) thereby, unlikely to be present in initiation complexes. When comparing TIE a3 with TIE a11, we could detect some initiation factors and translation-related proteins specific for a3 like eIF2D, eIF1A, eIF1A-X, eIF5B, LARS, QARS, DHX21 and HBS1L (**Fig. 6**). These results show a variation in the translation factors involved which hints two distinct TIE-mediated inhibitions. Similarly, we purified programmed pre-initiation complexes blocked by GMP-PNP (**Fig. S6**). By comparing a3 to either β-globin or a11 mRNAs, we found again an enrichment of eIF2D as a3-specific factor. Interestingly, we also found an enrichment of PKR protein (also called eIF2AK2) for both a3 and a11 (**Fig. S6**). PKR is known to bind double-stranded RNA during viral infection which mediates its auto-activation and induces the phosphorylation of eIF2α subunit (Adomavicius et al., 2019). This leads to the inhibition of mRNA translation. The enrichment of both eIF2D and PKR for a3 raised the question of whether eIF2D is indeed mediating the translation of a3 uORF by an alternative mechanism. To address this hypothesis and the involvement of eIF2D, we conducted a set of *in vivo* experiments.

**Figure 6:**
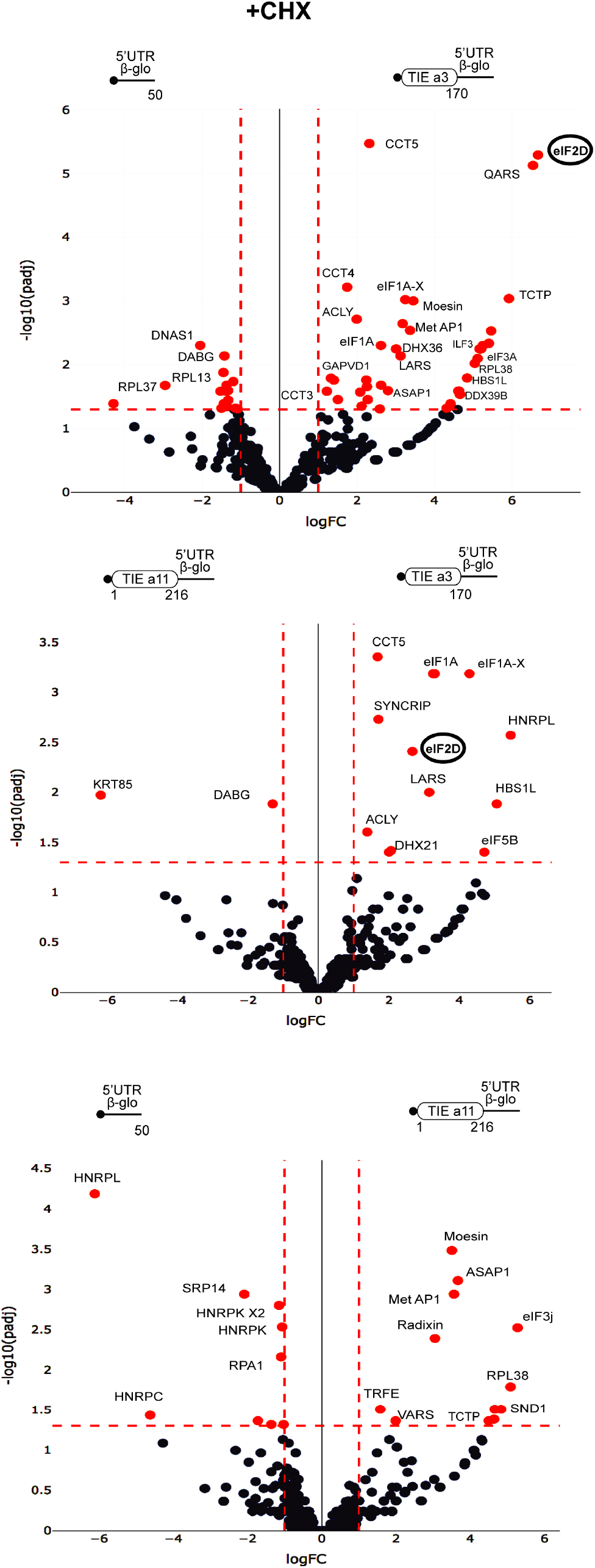
distinct profiles for factors involved in TIE-mediated inhibition blocked with cycloheximide. Mass spectrometry analysis of cycloheximide-blocked translation initiation complexes and on three transcripts: TIE a3, TIE a11 placed upstream of 5’UTR of β-globin and the 5’UTR of β-globin (control). Graphical representation of proteomics data: protein log_2_ spectral count fold changes (on the x-axis) and the corresponding adjusted log_10_ P-values (on the y-axis) are plotted in a pair-wise volcano plot. The significance thresholds are represented by a horizontal dashed line (P-value = 1.25, negative-binomial test with Benjamini–Hochberg adjustment) and two vertical dashed lines (−1.0-fold on the left and +1.0-fold on the right). Data points in the upper left and upper right quadrants indicate significant negative and positive changes in protein abundance. Protein names are labelled next to the off-centred spots and they are depicted according to the following color code: red spots are significant hits and black are non-significant with <10 spectra. Data points are plotted based on the average spectral counts from triplicate analysis. Three profiles were produced by comparing the proteomics of two transcripts.

### Translation inhibition by TIE a3 requires eIF2D

After specifically identifying eIF2D by Mass Spectrometry analysis for TIE a3-ribosomal complex, we were interested in getting more insights into how this factor might be involved. Previous studies have shown that eIF2D is a non-canonical translation initiation factor which delivers tRNA to the P-site of the ribosome in a GTP-independent manner (Dmitriev et al., 2010). It has been shown to be involved in the initiation step on specific mRNAs (Dmitriev et al., 2010; Vaidya et al., 2017; Vasudevan et al., 2020), reinitiation on main ORF (Ahmed et al., 2018; Weisser et al., 2017). Additionally, other studies showed that eIF2D is required for recycling after translation termination (Skabkin et al., 2010, 2013; Young et al., 2018). To confirm the involvement of eIF2D in TIE a3-mediated inhibition mechanism, we performed a co-transfection assay in HEK293T cell line using siRNA against either eIF2D, eIF4E or non-target pool of siRNAs together with a reporter plasmid harbouring TIE a3. As a control, we used reporter plasmids with TIE a11 and the same construct without TIE. After 48 hours incubation, cells were lysed for western blot analysis and luciferase assay **(Fig. 7A)**. Luciferase values were normalized to the reporter without TIE. We have previously shown that a3 and a11-mediated translation inhibition is achieved through cap-dependent translation of TIE uORFs. As expected, silencing of eIF4E (at efficiency of 74% and 76% respectively), which will significantly affect cap-dependent translation, drastically impairs a3- and a11-mediated translation inhibition. Interestingly, silencing eIF2D at an efficiency of 76%, also drastically abolishes a3-mediated inhibition and luciferase expression is restored to 85% **(Fig. 7A)**. On the contrary, the silencing of eIF2D at efficiency of 81% has no significant effect on a11-mediated inhibition. Altogether, this confirms the requirement of eIF2D for efficient a3 inhibition but not for a11. This conclusion is in good agreement with our previous Mass Spectrometry analysis of a3-ribosomal complexes blocked by translation inhibitors, which show the specific presence of eIF2D in the initiation complexes programmed with TIE-a3 **(Fig. 6 & S6)**. The fact that we found eIF2D in pre-initiation complexes suggests that eIF2D is involved in the initiation of a3 uORF. Since a3 uORF is long and extends downstream the AUG start codon of the main CDS in native Hoxa3 mRNA, which means that the a3 uORF partially overlaps the main CDS, a mechanism using a reinitiation event after a3 uORF translation is impossible. Therefore, we rather favour a model in which eIF2D is required for a3 initiation. Previous studies have shown that an A-motif upstream of an uAUG has been shown to be important for proper eIF2D recruitment (Dmitriev et al., 2010). Interestingly, a closer look at the sequences in the TIE a3 uORF revealed A-rich motifs upstream and downstream of uAUG111. We tested the implication of both A-motifs, AAAA107 upstream of the AUG and AAAAA147 downstream of the AUG, in TIE-a3 mediated inhibition (**Fig. 7B**). In order to avoid any side effect due to mutation in the sequence context around the uAUG, we kept an optimal Kozak sequence and mutated the As at position 107-110 to GGCC thereby keeping a purine residue at −3 position. The second A-motif downstream of AUG was similarly mutated to GGCCC. Interestingly, the mutation of upstream A-motif had a 2-fold reduction effect on translation inhibition of TIE a3. In contrast, mutation of the downstream A-motif does not affect TIE a3 inhibition (**Fig. 7B**). Additionally, we confirmed the implication of upstream A-motif *in vivo* with HEK293T cell line using plasmids with wild type a3 and the mutant of AAAA107 into GGCC (**Fig. 7B**). The AAAA107 mutant reduces the inhibitory efficiency of TIE a3 by 2.5 times compared to wild type a3. Therefore, we show that the A-motif upstream of the AUG is critical for translation inhibition and most probably because it is required for eIF2D recruitment.

**Figure 7:**
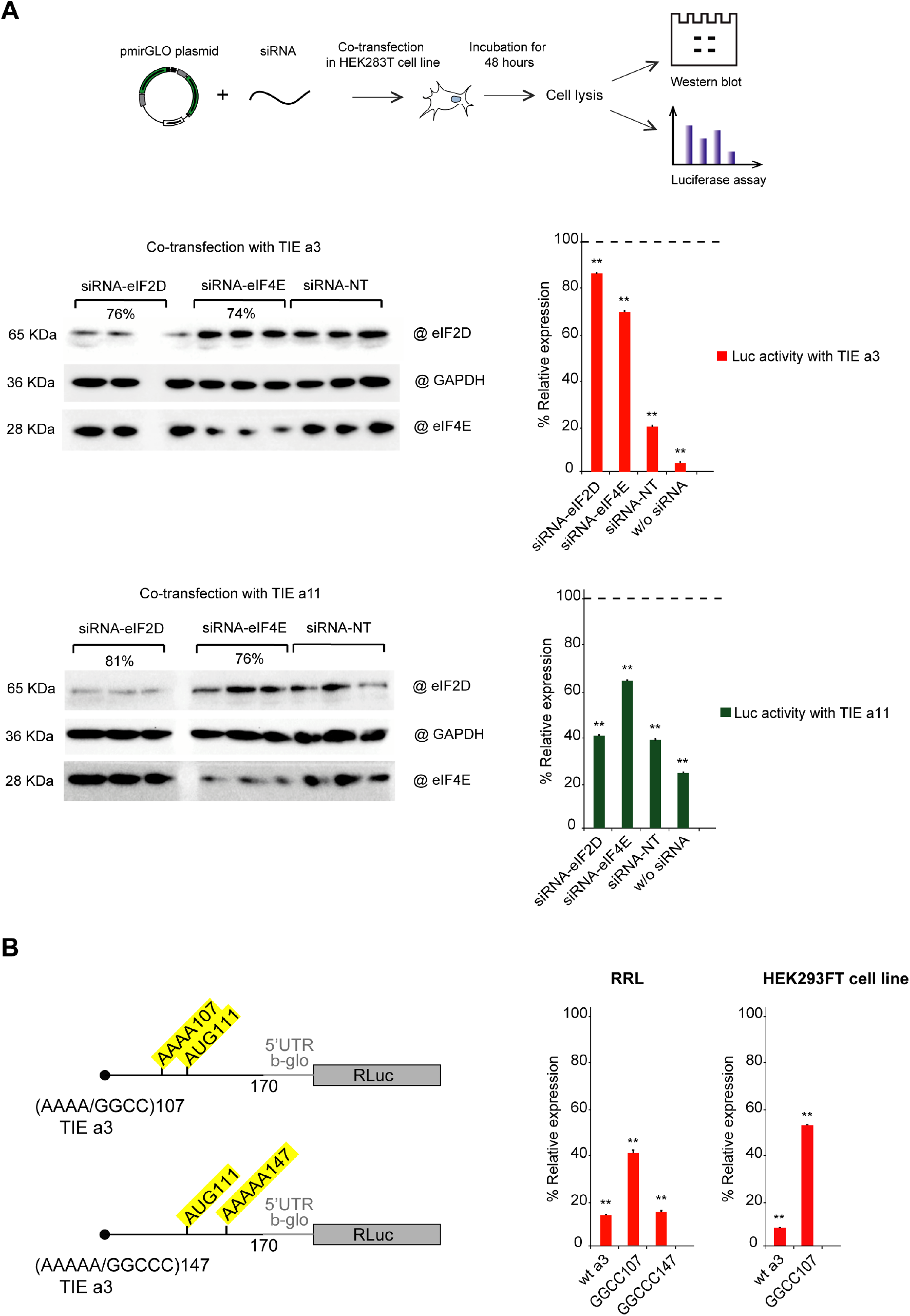
Co-transfection assay of TIE plasmids and siRNA against eIF2D confirms its implication in a3-mediated inhibition. Mutational analysis of A-motif sequences in TIE a3 shows a requirement for an upstream A-motif for efficient inhibition. **(A)** Co-transfection assay was performed using pmirGLO plasmids with TIE a3, TIE a11 and w/o TIE with siRNA against eIF2D, eIF4E and non-target pool of siRNA in HEK293 cell line. Plates were incubated for 48 hours followed by cell lysis. Cell lysate was analysed by western blot and luciferase assay. Luciferase activity was normalized compared to control (w/o TIE). **p<0.01 (t-test as compared to construct w/o TIE). n=3. Experiments were performed in triplicates. Efficiency of silencing of each protein was quantified by western blot analysis for TIE a3 and TIE a11 samples **(B)** Two sets of mutations were performed on distinct A-motif sequences in TIE a3. The first mutation is AAAA/GGCC107 and the second mutation AAAAA/GGCCC147. The transcripts were *in vitro* translated in RRL with TIE a3 Wt and control (w/o TIE). Results were confirmed *in vivo* in HEK293FT cell line. Luciferase activities were normalized to control (w/o TIE). **p<0.01 (t-test as compared to construct w/o TIE). n=3. Experiments were performed in triplicates.

### TIE a3 mediated 80S formation is cycloheximide-resistant

To further characterize the molecular mechanism of TIE a3-mediated inhibition, we performed sucrose gradient analysis with TIE a3 in the presence of different specific translation inhibitors (**Fig. 8**). Using GMP-PNP, a non-hydrolysable analogue of GTP that prevents subunit joining, a canonical 48S ribosome accumulates as expected on both TIE a3 and the control (Eliseev et al., 2018) (**Fig. 8**). With cycloheximide, a translocation inhibitor that binds the E-site of the ribosome (Schneider-Poetsch et al., 2010), we observe the accumulation of a homogeneous 80S with the control blocked on the initiating codon. Surprisingly, with a3, we obtained an unusual profile with an 80S complexes and complexes heavier complexes than normal 80S complexes (indicated with an arrow). The nature of these complexes is unknown but we speculate that they correspond to multiple stalled ribosomes on TIE a3 (**Fig. 8**). Interestingly, with geneticin, an A-site inhibitor that prevents 80S ribosome translocation (Vicens and Westhof, 2003), we observe the expected accumulation of a homogeneous 80S complex for both TIE a3 and control mRNAs (**Fig. 8**). This suggests that cycloheximide blockage is partially inefficient with TIE a3. One explanation of the heavier complexes observed with a3 in the presence of cycloheximide could be that multiple ribosomes are assembled and blocked on the two uAUGs and the main AUG **(Fig. 8)**. However, this possibility is ruled out by our previous toe printing analysis on TIE a3 mRNA that shows that a single toe print on uAUG111 is detected in the presence of cylcoheximide **(Fig. 3B)**. Moreover, we observe the expected 80S complex with geneticin confirming that there is only one initiation site on this mRNA. Therefore, another explanation would be that we are indeed observing translating ribosomes that partially escape blockage by cycloheximide. Cycloheximide binds in the E-site of the ribosome (Garreau de Loubresse et al., 2014). Interestingly, cryo-EM structures have shown that eIF2D also binds in the E-site. The presence of eIF2D in the pre-initiation complex might overlap the cylcoheximide binding site and thereby prevent at least partially the access of cycloheximide to its binding site on the ribosome (Weisser et al., 2017). Although preliminary, this explanation could bring another argument in favour of the implication of eIF2D in TIE a3-mediated inhibition.

**Figure 8:**
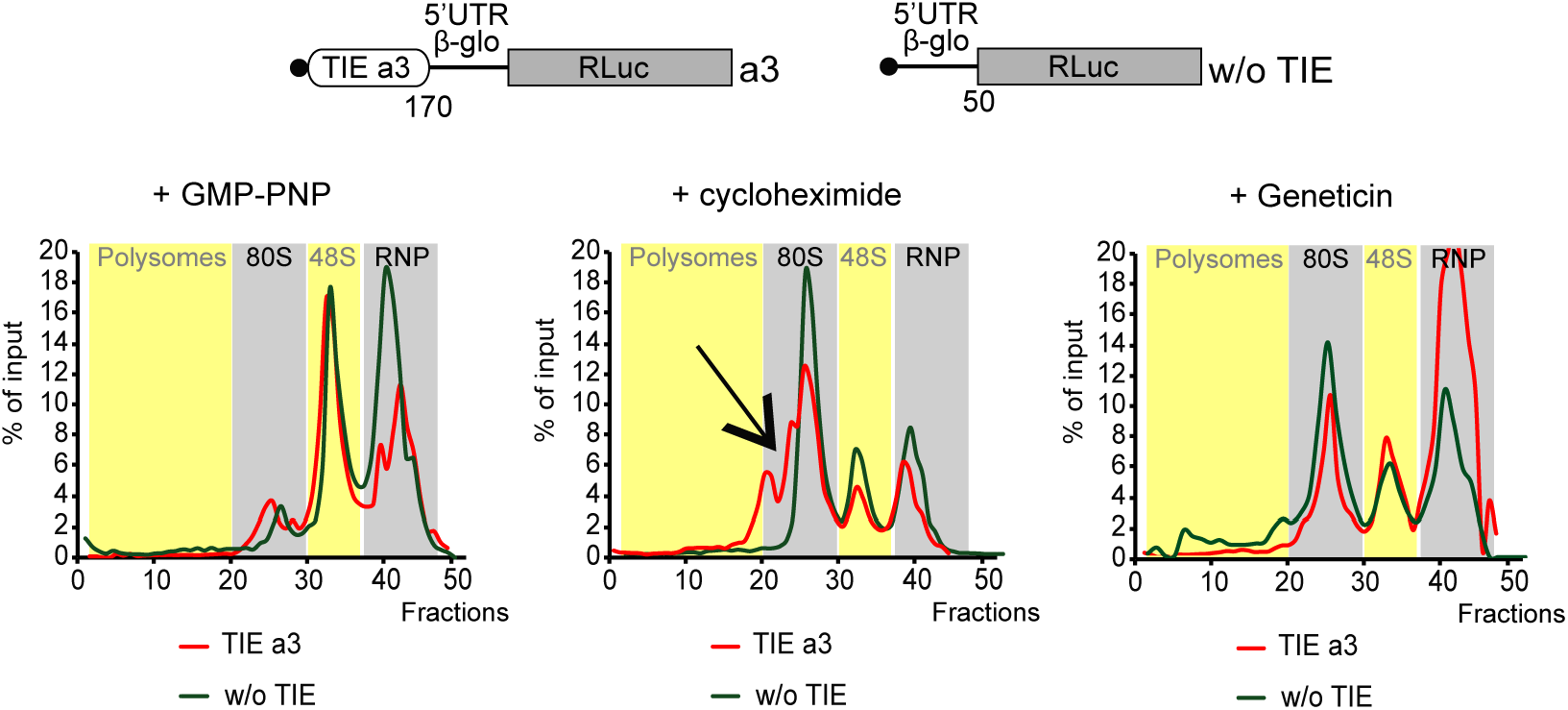
sucrose gradient analysis of TIE a3 with different inhibitors shows an atypical profile with cycloheximide. Ribosomal pre-initiation complexes were assembled and analysed on 7-47% sucrose gradient with radioactively capped TIE a3 and control (w/o TIE) in the presence of: GMP-PNP and translocation inhibitors cycloheximide and geneticin.

## Discussion

Our study has shown that two HoxA mRNAs, a3 and a11, are regulated by different mechanisms to ensure the inhibition of cap-dependent translation and allowed us to propose two distinct models for their mode of action (**Fig. 9**). First, we have shown that TIE elements can function *in vitro* using cell-free translation extracts. We then confirmed the results obtained with these extracts using *in vivo* assays in several cell lines. Our findings suggest that a3 inhibits translation by a uORF which is translated through the whole 5’UTR of Hox a3 mRNA producing a small protein of size 9 KDa. Interestingly, the alignment of Hox a3 TIE element shows a conservation of the uAUG111 (highlighted in red box) among different species. In contrast to the localisation of the uAUG that is highly conserved, the coding sequence of the uORF is not conserved among species (**Fig. S7A)**. Indeed, uORFs have been recognized as regulators of translation for number of cellular mRNAs (Barbosa et al., 2013). For instance, four uORFs in the 5’ leader of *GCN4* mRNA restrict the flow of scanning ribosomes from the cap site to the GCN4 initiation codon (Dever et al., 2016; Hinnebusch, 1993). The uORF in AdoMetDC mRNA generates a nascent hexapeptide that interacts with its translating ribosome to suppress translation of AdoMetDC RNA in a cell-specific manner (Uchiyama-Kadokura et al., 2014). Interestingly, our data indicates that eIF2D is required in TIE a3 mode of action. A Recent study has shown that ATF4 mRNA translation is induced by eIF2D and its homologue DENR during integrated stress response (Vasudevan et al., 2020). In this case, eIF2D requires its RNA binding motif to mediate translation of ATF4 mRNA through its 5’ leader sequence consisting of multiple uORFs (Vasudevan et al., 2020). Moreover, it has been shown that eIF2α-phosphorylation during stages of embryonic development promotes translation from uORFs (Friend et al., 2015). Therefore, canonical cap-dependent translation initiation with eIF2 is not possible during embryonic development. The cap-dependent translation initiation of uORF from TIE a3 might use eIF2D as an alternative to replace inactive phosphorylated eIF2 to promote uORF translation. So far, the only *cis*-acting sequence that has been clearly defined on an mRNA for specific eIF2D recruitment is an A-rich motif upstream of the start codon (Dmitriev et al., 2010). We showed that TIE a3 contains such an A-motif upstream of the uORF that is critical for TIE function (**Fig. 7**). Other reports showed that eIF2D would form initiation complexes on leaderless and A-rich 5’UTRs (Akulich et al., 2016; Dmitriev et al., 2010). In TIE a3, mass spectrometry analysis enabled us to demonstrate that eIF2D is present only in pre-initiation complexes programmed with TIE a3. Interestingly, we also showed that cycloheximide does not fully inhibit TIE a3 uORF translation as expected which suggests that eIF2D might at least partially interfere with cylcoheximide binding in the ribosomal E-site. In fact, other examples of partial cycloheximide resistance of specific mRNAs have been recently described (Kearse et al., 2019). In these cases, cycloheximide resistance occurs due to queuing of scanning preinitiation complexes in response to slowly elongating ribosomes from non-AUG codons. Regarding TIE a11, the ribosome recognizes a combination of two *cis-acting* elements in the 5’UTR (**Fig. 9**). (i) A start-stop codon combination located at positions 84-89 and (ii) a long highly stable GC-rich helical structure (SL) located at +20 downstream of the uAUG (by convention, the A from the AUG being +1). These two elements act in synergy to promote the stalling of an 80S complex upstream of the SL. This is a reminiscent of a similar mechanism that has been described in the *Arabidopsis thaliana* NIP5.1 5’UTR mRNA that contains an AUG-stop that regulates translation of the main ORF through a ribosome stalling mechanism and mRNA degradation (Tanaka et al., 2016). This mechanism requires only the start-stop codons. In the case of TIE a11, an additional *cis*-element is required for ribosome stalling, namely the stable SL that is present downstream of the AUG-stop. A ribosome that is stalled with a stop codon in the A site, in other words an empty A site without any tRNA, is usually the signal for recruitment of the release factors to dissociate the ribosomal subunits from the mRNA. With our cell-free translation assay and sucrose gradient analysis, we showed that the stalled 80S programmed with TIE a11 is very stable and does not dissociate **(Fig. 5C)**. A possible explanation for the stability of this complex could be that the SL blocks the access of the release factors to the empty A site thereby preventing ribosome dissociation (Brown et al., 2015). Interestingly, the alignment of TIE a11 among different species shows that the start-stop combination (highlighted in red) lacks conservation (**Fig. S7B).** Some species possess a substitution of the start codon AUG to AUG-like codons such GUG or a mutation of stop codon that leads to a longer uORF. In contrast, the SL (104-154) (highlighted in green) remains highly conserved suggesting a functional significance (**Fig. S7B**). Accordingly, we have shown that the sole SL is strong enough to impede scanning by the pre-initiation complex. Indeed, secondary structures in the 5’UTR have been shown to inhibit translation like the case of a conserved stem loop structure in the 5’UTR of TGF-β1 mRNA (Jenkins et al., 2010). Therefore, in various species, TIE a11 might use different combinations of *cis*acting elements in order to block cap-dependent translation, the common *cis*-acting element between all species being the SL that is conserved.

**Figure 9:**
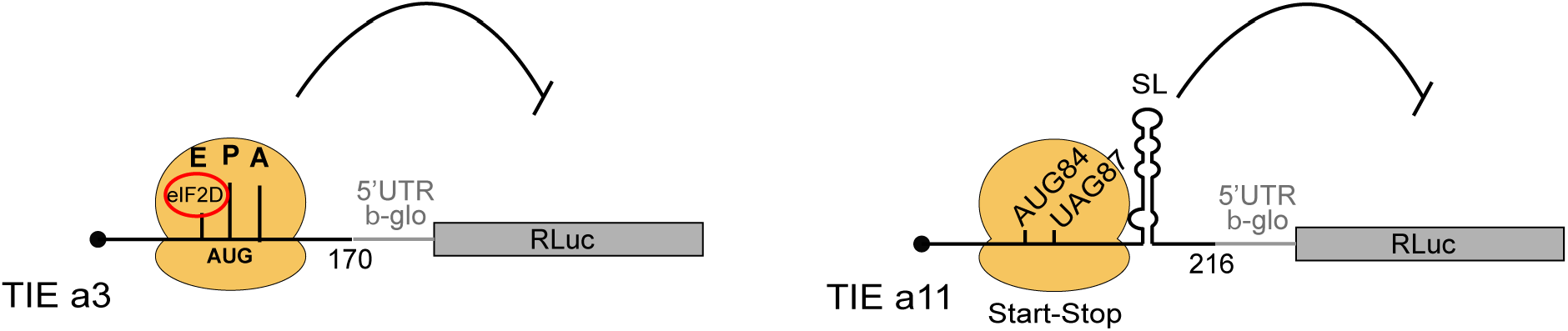
two distinct models for translational inhibition by TIE a3 and TIE a11. A model for TIE a3 suggests a ribosomal assembly on the uAUG111 with a requirement of eIF2D initiation factor. The model for TIE a11 suggests a stalled 80S ribosome on AUG-stop codons combination upstream of a highly stable structure.

As previously described, the IRES in Hox a11 mRNA requires the presence of the ribosomal protein RpL38 while that in Hox a3 mRNA is RpL38-independent (Kondrashov et al., 2011).

Our probing experiments revealed that the folding of both a3 and a11 TIE elements is independent of the presence of IRES suggesting that their mode of action does not depend on the IRES. TIE may have evolved in such a way to favour the translation from the downstream IRES hence justifying why there is variation in terms of sequence and mechanism but same inhibitory effect. This unique combination of an inhibitor of canonical translation mechanism and the activator of specialized translation sets an interesting point on how the 5’UTR elements confer ribosome specificity to translation (Xue et al., 2015). Importantly, the acquisition of these TIE elements in subsets of Hox mRNAs enables an additional layer of regulatory control between the canonical translation and the IRES-dependent one. One intriguing study would be to determine how other TIE elements a4 and a9 inhibit translation and whether there are common functional features amongst all Hox mRNAs. Beyond Hox mRNAs, our data on eIF2D suggests a specific role in translation initiation and it would be interesting to decipher its precise role at the molecular level. More precisely, determining how uORF length, codon composition and consensus sequences may influence the role of eIF2D in the initiation process on uORFs. Future studies will be required to fully understand the role of eIF2D in translation initiation of specific mRNAs.

## Acknowledgments

This work was funded by ‘Agence Nationale pour la Recherche’ (Ribofluidix, ANR-17-CE12-0025-01, by University of Strasbourg and by the ‘Centre National de la Recherche Scientifique’. We would like to thank Dr. Maria Barna for sharing sequences of Hox a3 and 11 transcripts. We are grateful to Dr. Christine Allmang, Hassan Hayek, Antonin Tidu, Aurélie Janvier, Aurélie Durand for technical assistance, Philippe Hamman, Johana Chicher, Lauriane Kuhn for Mass spectrometry analysis, Dr. Mireille Baltzinger and Dr. Pascale Romby for support and useful discussions on the project. We would also like to thank Dr. Sebastien Pfeffer for the pmirGLO plasmid.

## Author contributions

FM, GE and FA designed the experimental strategy, FA performed all the experiments with the technical assistance of LS. FA, GE and FM interpreted the results and wrote the manuscript.

## Supplemental Figure Legends

**Supplemental figure S1:**
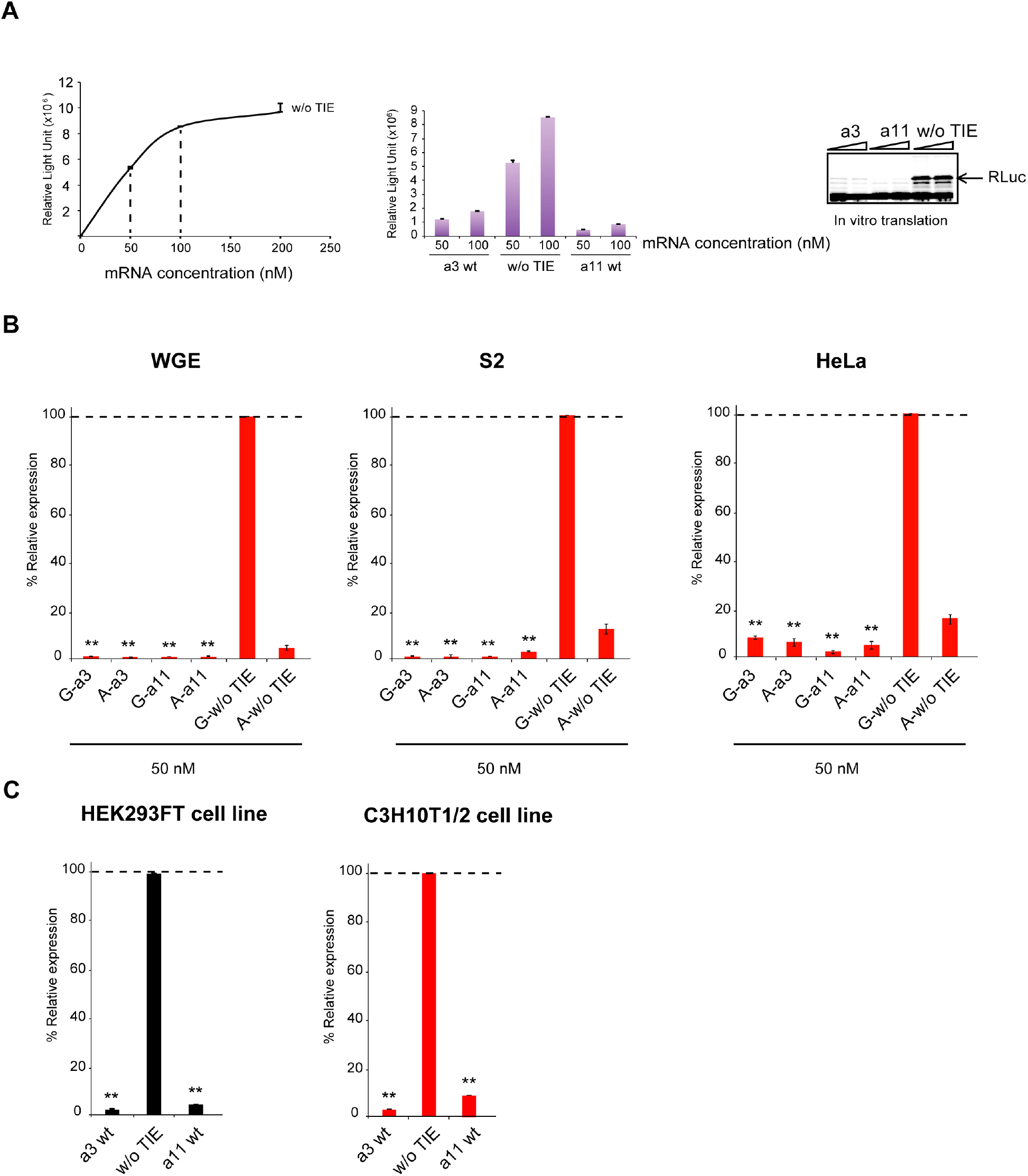
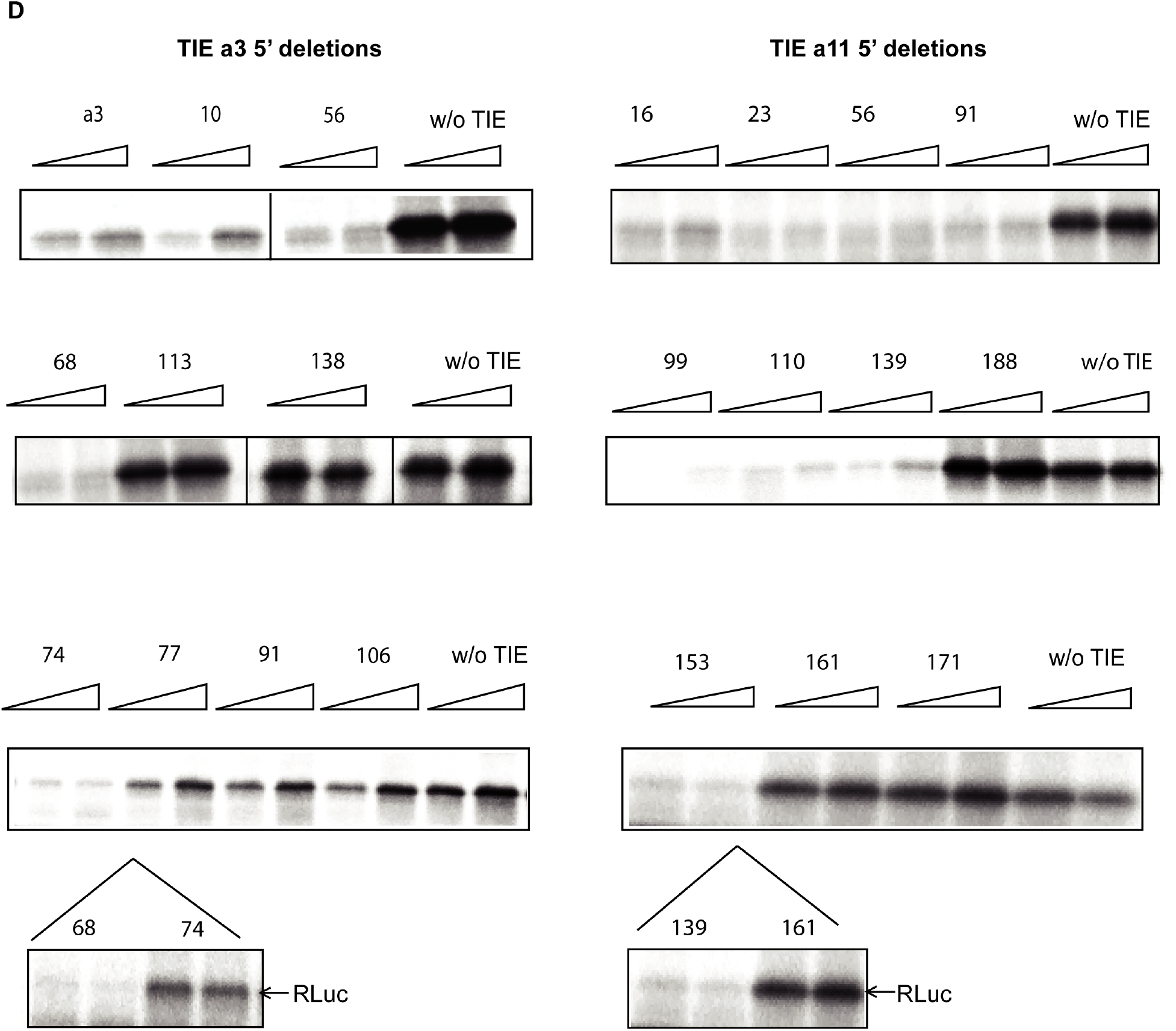
TIE-mediated inhibition recapitulated in different *in vitro* and *in vivo* systems. **(A) (To the left)** The intensity of light emitted by Renilla Luciferase protein as a function of control mRNA (w/o TIE) concentration (nM). Emission of light was measured under subsaturating conditions (50-100 nM mRNA concentration). **(Middle panel)** The intensity of light emission by RLuc protein relative to different concentrations (50 nM and 100 nM respectively) of tested mRNA samples TIE a3, TIE a11 and control (w/o TIE). **(To the right)** Analysis of translation products after *in vitro* translation in RRL on 10% SDS PAGE to monitor RLuc expression. Visualization of protein bands was achieved by incorporation of radiolabelled ^35^S-methionine, which are detected by autoradiography. **(B)** *In vitro* translation of TIE a3 and TIE a11 transcripts in the presence of m^7^G_ppp_G cap or a non-functional analog A_ppp_G. Three *in vitro* systems were used: Wheat Germ Extract (WGE), drosophila embryonic cell extract (S2) and HeLa cell extract. All mRNAs were translated *in vitro* at 50 nM concentrations and RLuc expression was normalized to control (w/o TIE) in each condition. **(C)** Transfection of reporter plasmids with TIE a3 or TIE a11 in two embryonic cell lines, kidney HEK293FT and mesenchymal C3H10T1/2 cell lines. Renilla luciferase expression was normalized to the control (w/o TIE). **p<0.01 (t-test as compared to construct w/o TIE). n=3. Experiments were performed in triplicates. **(D)** SDS PAGE of Renilla expression with 5’ deletions in TIE a3 and TIE a11. The gels at the bottom refer to the minimal functional region compared to longest construct that is inactivated by 5’ deletions.

**Supplemental figure S2:**
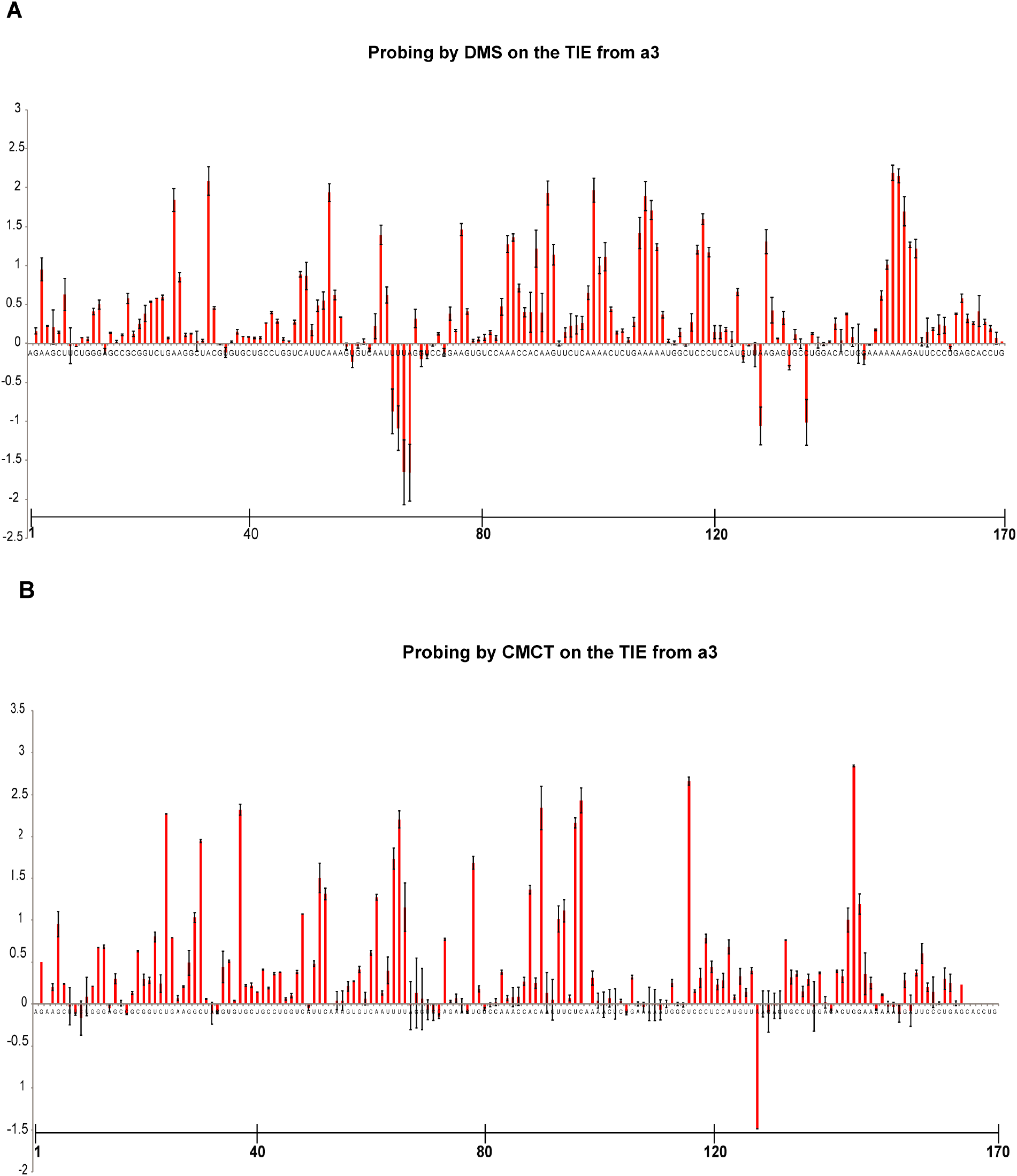
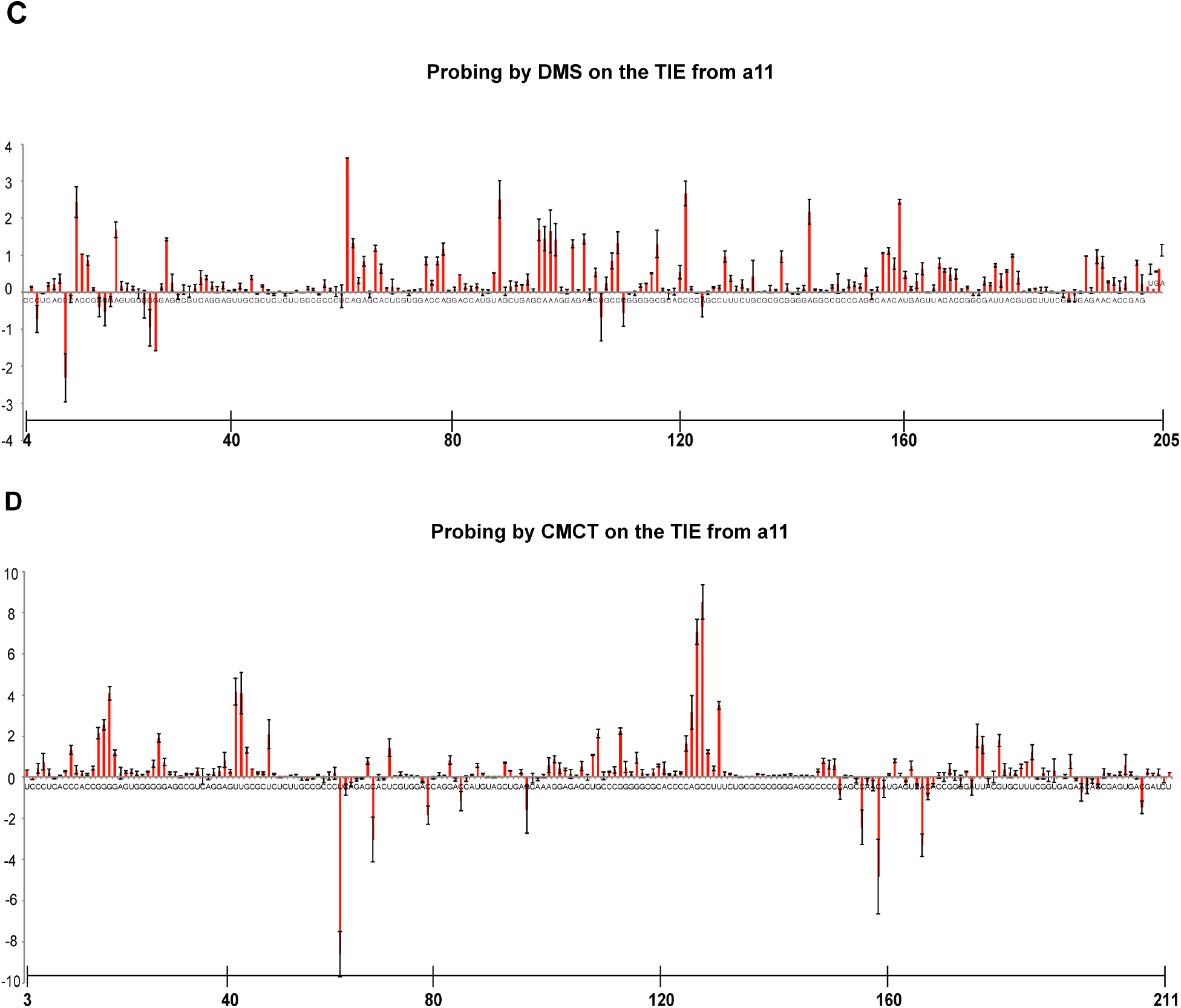
average of reactivities of DMS and CMCT for TIE a3 and TIE a11. Probing experiments by DMS and CMCT were performed in triplicates. Figures S2A and S2B represent averages of reactivities of DMS and CMCT for TIE a3 respectively. Figures S2C and S2D represent averages of reactivities of DMS and CMCT for TIE a11 respectively. A scale of 40 nucleotides range was included to determine position of every nucleotide. Standard deviations are shown.

**Supplemental figure S3:**
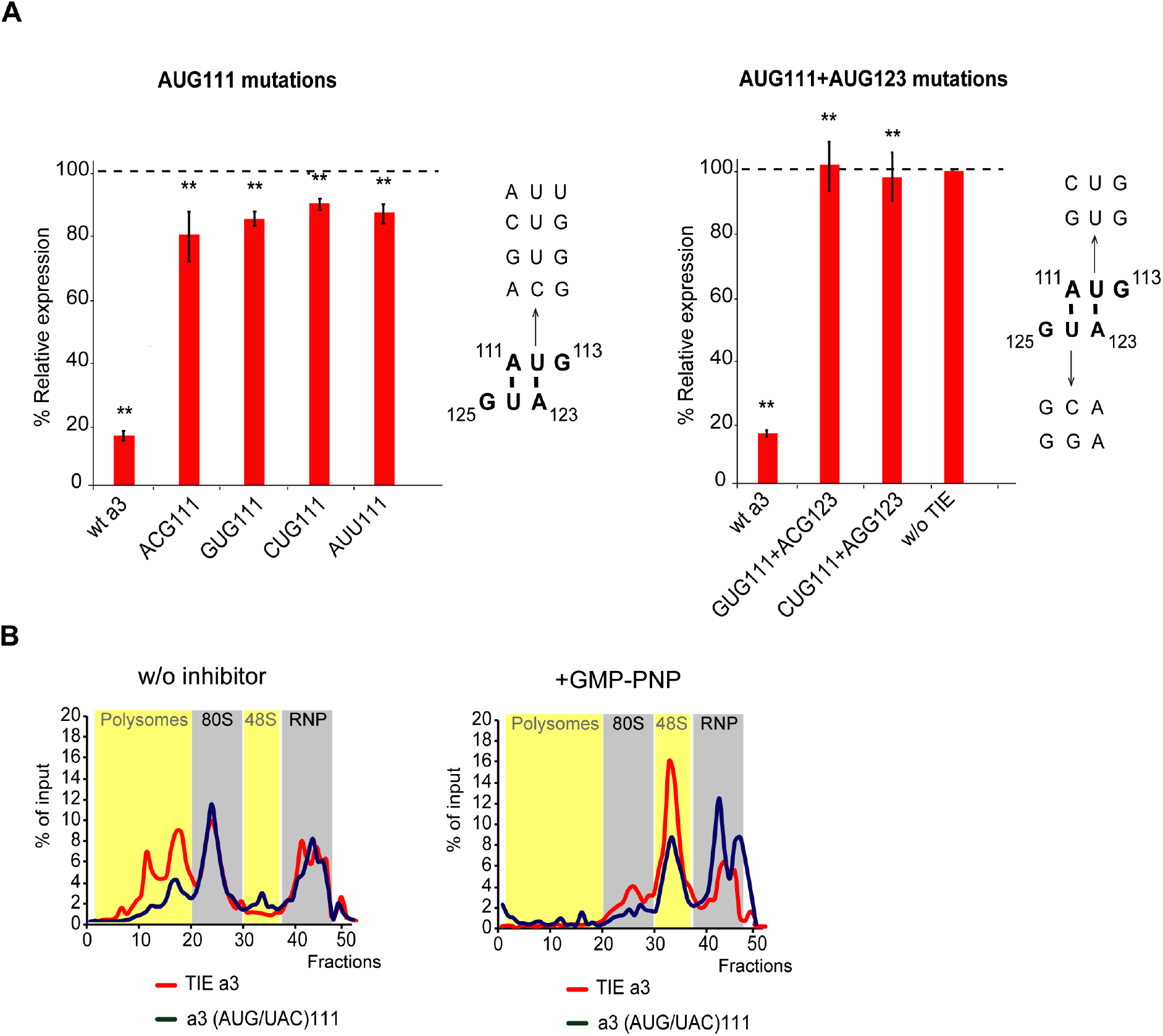
upstream AUG111 in TIE a3 is essential for inhibition. **(A) (To the left)** AUG-like mutations (AUU, GUG, CUG and ACG) were tested compared to wt a3. (**To the right)** Representation of uAUG111 context in the secondary structure of TIE a3 is shown. A111 and U112 base pair with the A123 and U124 of the second AUG123. Mutants of the uAUG into CUG and GUG with the corresponding compensatory mutagenesis were tested. RLuc expression with different transcripts was normalized to control (w/o TIE). **p<0.01 (t-test as compared to construct w/o TIE). n=3. Experiments were performed in triplicates. **(B)** Sucrose gradient analysis of TIE a3 and mutant of upstream (AUG/UAC)111. Ribosomal pre-initiation complexes were assembled and analysed on 7-47% sucrose gradient with radioactive m^7^G-capped TIE a3 and the mutants of upstream (AUG/UAC)111 in the absence (upper panel) or in the presence of GMP-PNP (lower panel). Heavy fractions correspond to polysomes while the lightest correspond to free RNPs.

**Supplemental figure S4:**
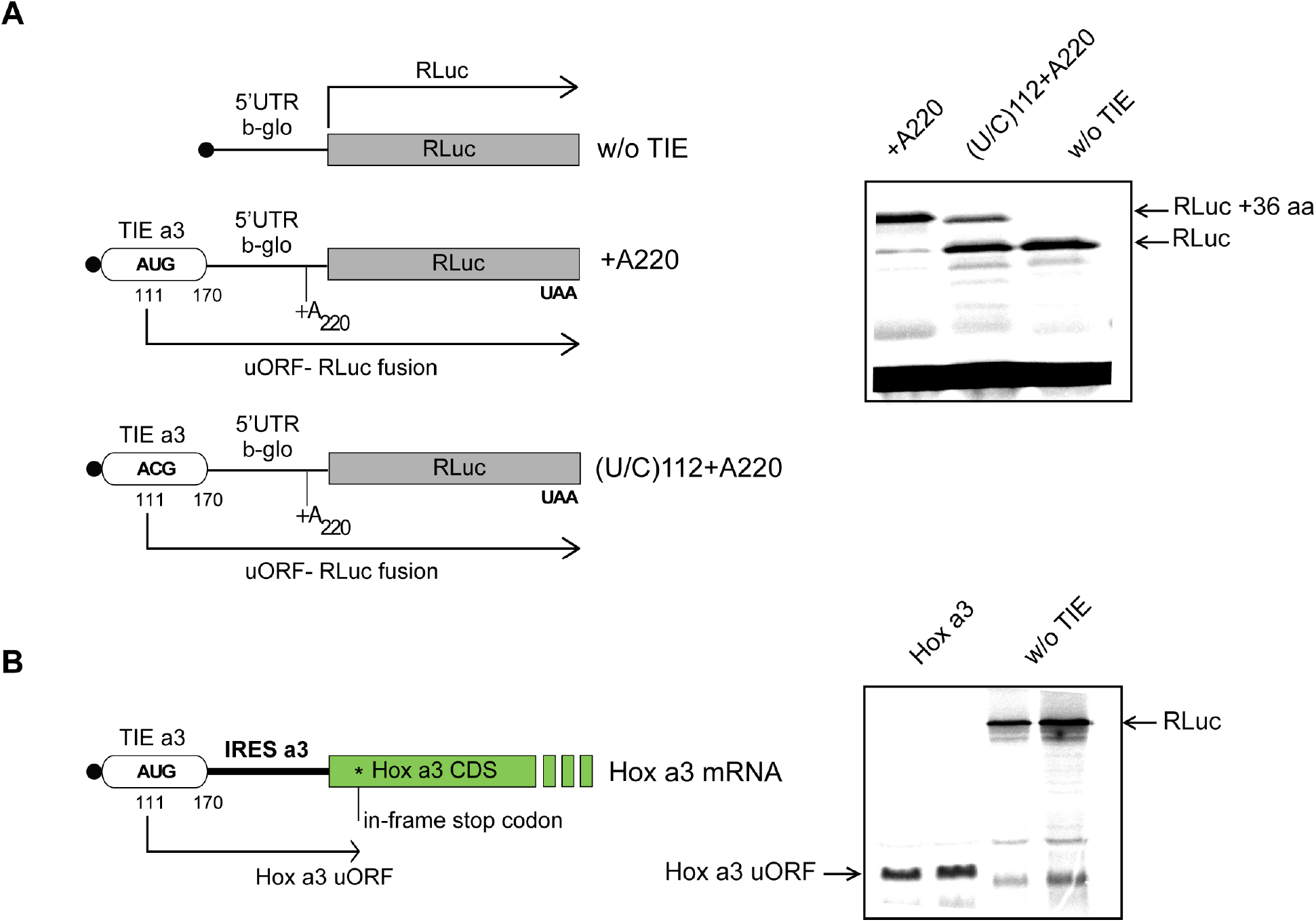
translation of TIE a3 uORF in different constructs. **(A)** A single nucleotide deletion of A220 in TIE a3 is performed to frameshift the uORF frame to the same as the main Rluc ORF. Another mutation was performed combining this deletion with uAUG mutation of (U/C)112. Transcripts were *in vitro* translated in RRL and products were loaded on 10% SDS PAGE. **(B)** Translation of TIE a3 uORF in Hox a3 mRNA. Hox a3 mRNA (5’UTR+CDS) was translated *in vitro* to check uORF translation in this context. Products were analysed on 15% SDS-PAGE.

**Supplemental figure S5:**
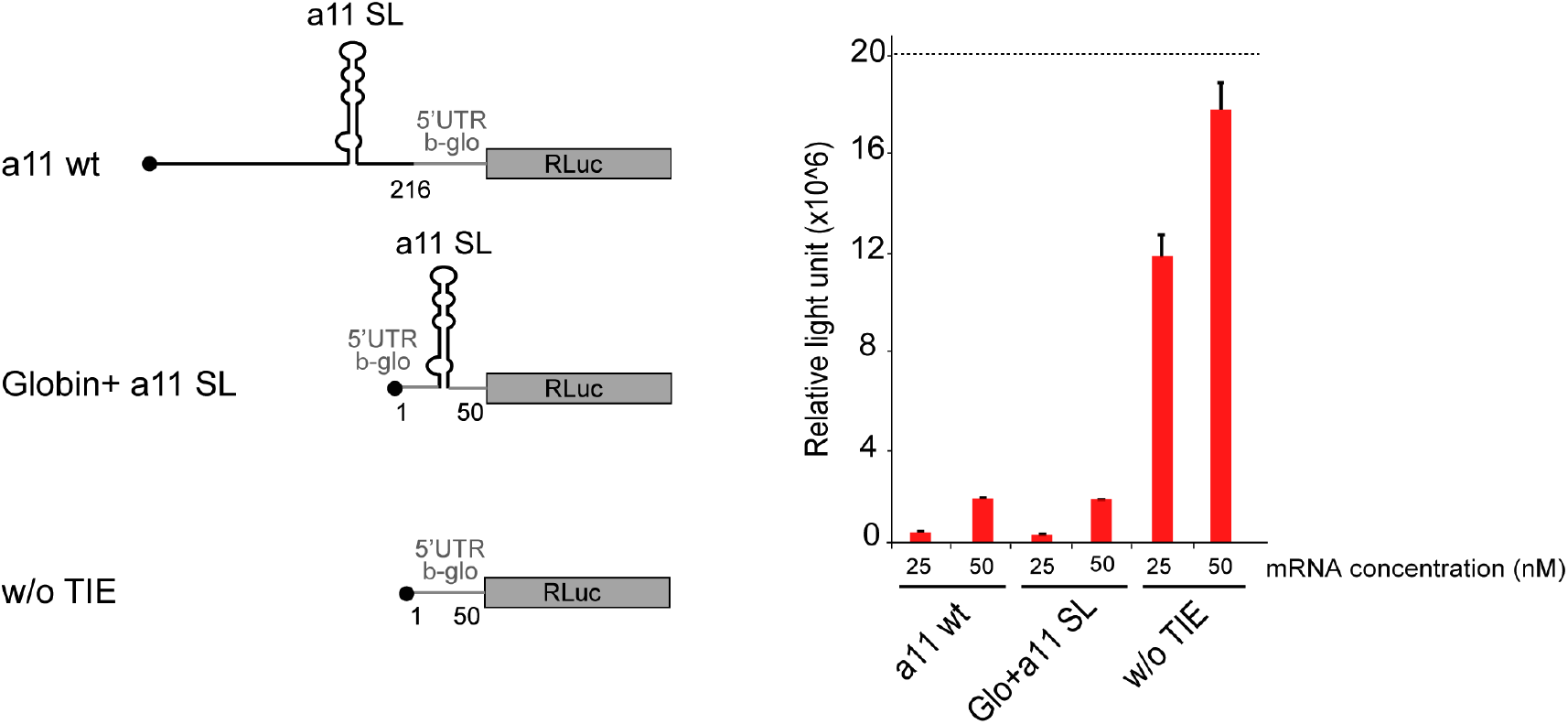
transplanting TIE a11 stem loop structure in globin 5’ UTR efficiently inhibits translation of RLuc mRNA. The stem loop structure of TIE a11 (104-154) was transplanted in the 5’UTR of β-globin with 25 nucleotides spanning from each extremity. The three shown transcripts were *in vitro* translated in RRL at two concentrations, 25 nM and 50 nM. Results are represented in relative light unit of Rluc expression.

**Supplemental figure S6:**
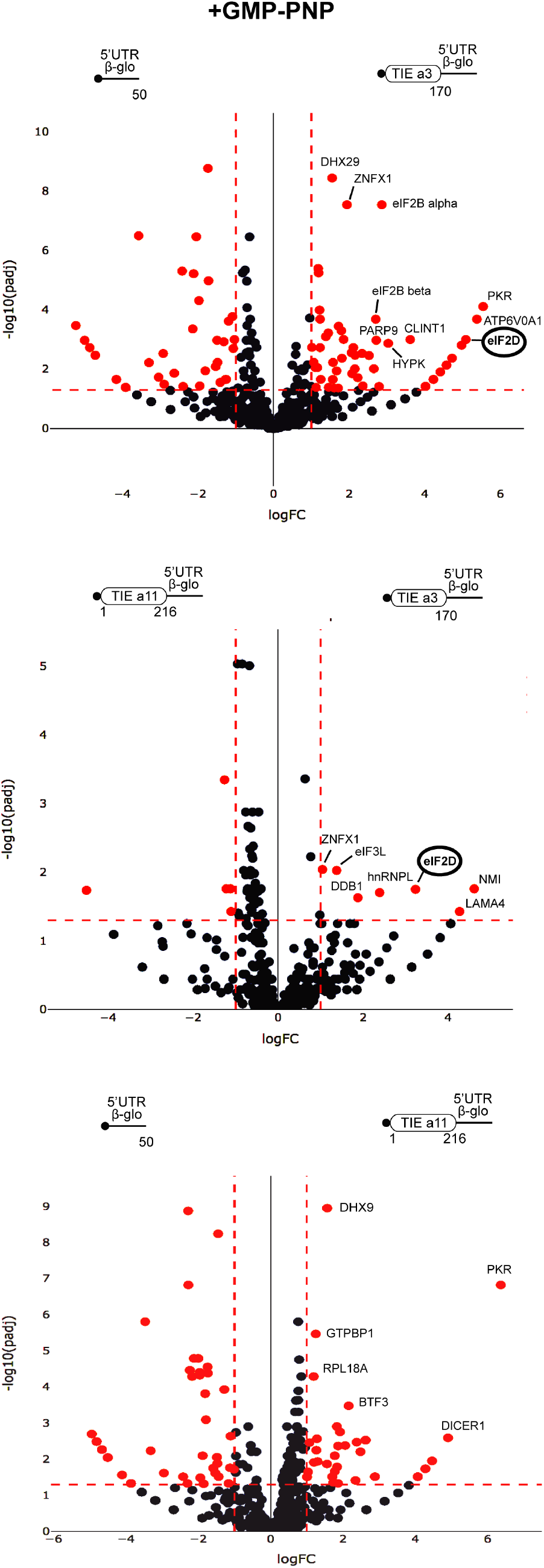
distinct profiles for factors involved in TIE-mediated inhibition blocked with GMP-PNP. Mass spectrometry analysis of GMP-PNP-blocked translation initiation complexes and on three transcripts: TIE a3, TIE a11 placed upstream of 5’UTR of β-globin and the 5’UTR of β-globin (control). Graphical representation of proteomics data: protein log_2_ spectral count fold changes (on the x-axis) and the corresponding adjusted log_10_ P-values (on the y-axis) are plotted in a pair-wise volcano plot. The significance thresholds are represented by a horizontal dashed line (P-value = 1.25, negative-binomial test with Benjamini–Hochberg adjustment) and two vertical dashed lines (−1.0-fold on the left and +1.0-fold on the right). Data points in the upper left and upper right quadrants indicate significant negative and positive changes in protein abundance. Protein names are labelled next to the off-centred spots and they are depicted according to the following color code: red spots are significant hits and black are non-significant with <10 spectra. Data points are plotted based on the average spectral counts from triplicate analysis. Three profiles were produced by comparing the proteomics of two transcripts.

**Supplemental figure S7:**
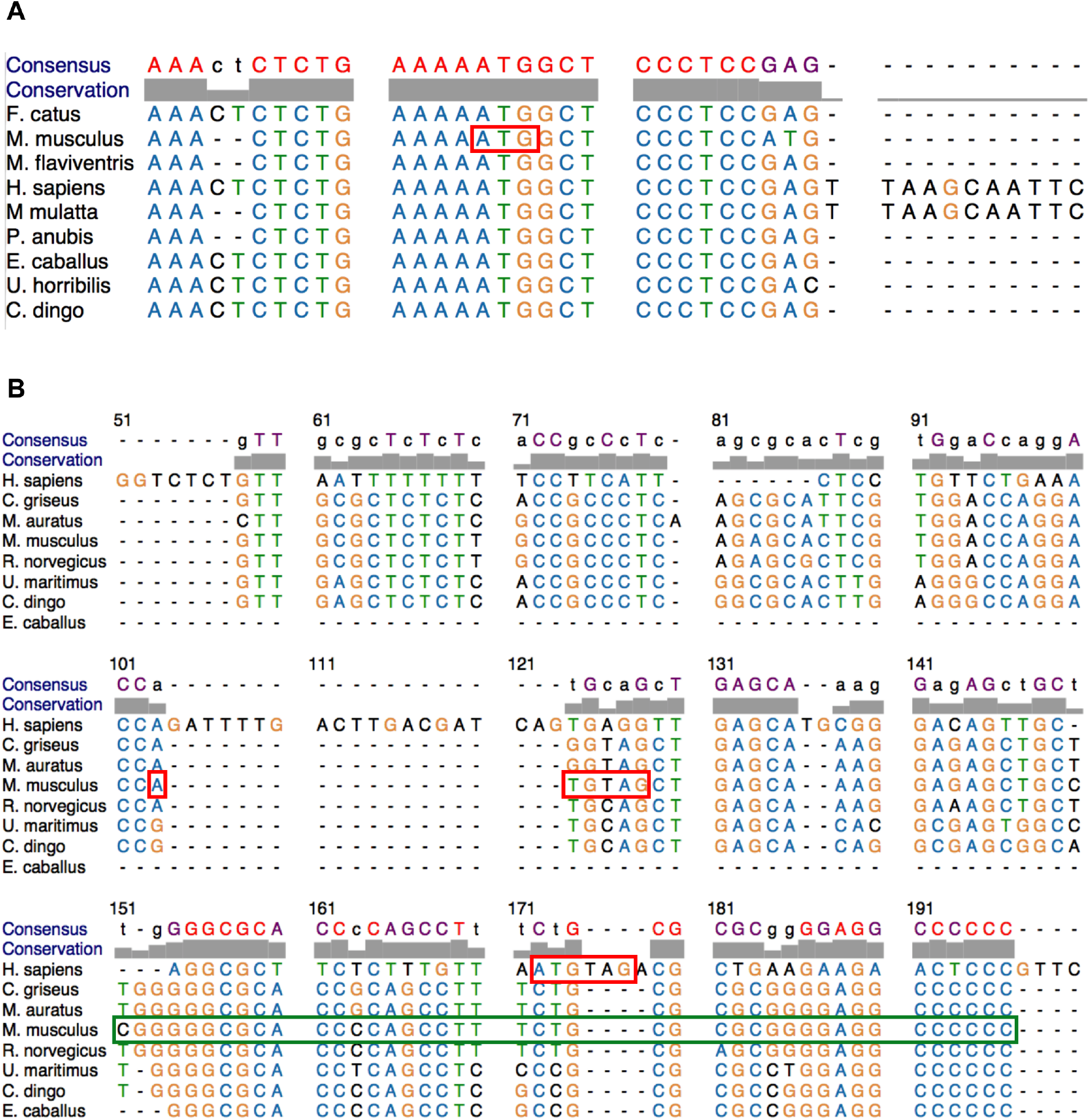
Alignment of TIE a3 and TIE a11 among different species shows variation in the conservation of the inhibitory elements. **(A)** Nucleotides 1-170 from the mouse TIE a3 sequence were aligned with different species. The position of uAUG in *Mus musculus* is highlighted in a red box. Accession numbers: *F. catus*: XR_002740526.1, *M. musculus*: NM_010452.3, *M. flaviventris*: XM_0279, *H. sapiens*: XM_011515343.3, *M. mulatta*: XM_028845931.1, *P. Anubis*: XM_017956197.2, *E. caballus*: XM_023639416.1, *U. horribilis*: XM_026508219.1, *C. dingo*: XM_025464436.1 **(B)** Nucleotides 1-216 from the mouse TIE a11 sequence were aligned with different species. The position of uAUG-UAG in *Mus Musculus* and the insertion of AUG-UAG in *Homo sapiens* are highlighted in red boxes. Accession numbers: *H. sapiens*: AF071164.1*, M. Mulata*: XM_015133828.2, *C. dingo*: XR_003143092.1, *C. griseus*: XM_027436289.1, *R. norvegicus*: XM_008762951.2, *E. caballus*: XM_023639423.1, *M auratus*: XM_00508585871.3, *M. musculus*: NM_010450.

